# Promoter-specific dynamics of TATA-binding protein association with the human genome

**DOI:** 10.1101/696369

**Authors:** Yuko Hasegawa, Jason D. Lieb, Kevin Struhl

## Abstract

Transcription factor binding to target sites *in vivo* is a dynamic process that involves cycles of association and dissociation, with individual proteins differing in their binding dynamics. The dynamics at individual sites on a genomic scale has been investigated in yeast cells, but comparable experiments have not been done in multicellular eukaryotes. Here, we describe a tamoxifen-inducible, time-course ChIP-seq approach to measure transcription factor binding dynamics at target sites throughout the human genome. As observed in yeast cells, the TATA-binding protein (TBP) typically displays rapid turnover at RNA polymerase (Pol) II-transcribed promoters, slow turnover at Pol III promoters, and very slow turnover at the Pol I promoter. Interestingly, turnover rates vary widely among Pol II promoters in a manner that does not correlate with the level of TBP occupancy. Human Pol II promoters with slow TBP dissociation preferentially contain a TATA consensus motif, support high transcriptional activity of downstream genes, and are linked with specific activators and chromatin remodelers. These properties of human promoters with slow TBP turnover differ from those of yeast promoters with slow turnover. These observations suggest that TBP binding dynamics differentially affect promoter function and gene expression, possibly at the level of transcriptional reinitiation/bursting.

## INTRODUCTION

Binding of transcription factors to specific genomic DNA sequences is required for accurate and regulated transcription by RNA polymerases. This ensures biologically appropriate levels of RNA transcripts for a wide variety of environmental and developmental conditions. Transcription factor binding *in vivo* is analyzed conventionally by chromatin immunoprecipitation (ChIP), which measures occupancy at target sites on a cell- and time-averaged basis (Struhl 2007). However, ChIP represents a static measurement that does not consider the dynamics of binding, namely the dissociation and reassociation of proteins with their target sites.

FRAP (Fluorescence Recovery after Photobleaching) experiments indicate that many transcription factors show highly dynamic binding with very rapid dissociation rates, while other proteins (e.g. histones) have much slower turnover (McNally et al. 2000; Houtsmuller 2005; Mueller et al. 2010). However, FRAP experiments typically measure the average dynamic properties of a given protein on all target sites, and it is not possible to distinguish unbound vs. DNA-bound proteins in the bleached area. Using live-cell imaging, binding dynamics at specific DNA sites can be visualized on artificially tandem arrays of binding sites (McNally et al. 2000) or at genomic regions consisting of naturally occurring repeats (Karpova et al. 2008). In yeast, binding dynamics on endogenous single-copy genes can be measured on an individual basis by using a quench flow apparatus to perform a formaldehyde time course on a sub-second scale (Poorey et al. 2013). However, none of these methods address whether binding dynamics are uniform or variable over the entire range of target sites.

Genome-scale, site-specific analysis of transcription factor binding dynamics has been performed in yeast using a competition-ChIP approach (Dion et al. 2007; van Werven et al. 2009; Lickwar et al. 2012). Expression of an epitope-tagged transcription factor is induced by the addition of galactose, and whole-genome ChIP measurements are made at various times after induction. The kinetics of binding by the induced protein (distinguished by its epitope tag) at each target site provides information on protein turnover that site. Analyses of yeast TATA-binding protein (TBP), Rap1, and histone H3 by competition ChIP reveal that binding dynamics are variable at their target sites in a manner that is poorly correlated with occupancy levels determined by conventional ChIP (Dion et al. 2007; van Werven et al. 2009; Lickwar et al. 2012). Comparable experiments have not been performed in any multicellular organism.

Here, we describe a tamoxifen-inducible, time-course ChIP-seq analysis that permits the measurement of transcription factor binding dynamics at target sites on a genome-wide scale in human cells. We apply this method to analyze the dynamics of the TATA-binding protein (TBP). *In vivo*, TBP is required for transcription from promoters mediated by all three nuclear RNA polymerases (Pol) (Cormack and Struhl 1992). These three classes of promoters are responsible for the synthesis of rRNA (Pol I), mRNA and other RNAs (Pol II), and tRNA and other RNAs (Pol III). In mammalian cells, TBP does not bind promoters on its own, but rather as part of multiprotein complexes that are specific for the three promoter classes (SL1 for Pol I, TFIID for Pol II, and TFIIIB for Pol III) (Sharp 1992; Hernandez 1993; Struhl 1994).

Here, we show that TBP has strikingly different binding dynamics at human Pol I, Pol II, and Pol III promoters, similar to what is observed in yeast cells (van Werven et al. 2009; Poorey et al. 2013). Binding dynamics vary considerably among Pol II promoters in a manner that is poorly correlated with TBP occupancy. Consensus TATA motifs and the association of specific activators and chromatin remodelers are determinants of TBP binding turnover rate, and high transcription of downstream genes is linked to promoters with slower TBP turnover rates. We suggest that differences in TBP binding dynamics may be related to differences in transcriptional reinitiation or bursting.

## RESULTS

### Measuring dynamics of transcription factor binding to target sites in human cells

The genome-scale assay involves the rapid induction of an epitope-tagged protein and performing ChIP-seq analysis at various times after induction (Fig. 1A). The binding kinetics of the epitope-tagged protein provides a measurement of the turnover of the untagged, endogenous protein previously bound to the same sites. Although this method doesn’t measure the absolute dissociation rate of the endogenous protein, it is suitable for measuring relative turnover rates at different loci. Specifically, the target sites are simultaneously assayed with the same mixture of epitope-tagged and untagged endogenous protein at each time point (Fig. 1A).

**Figure 1.**
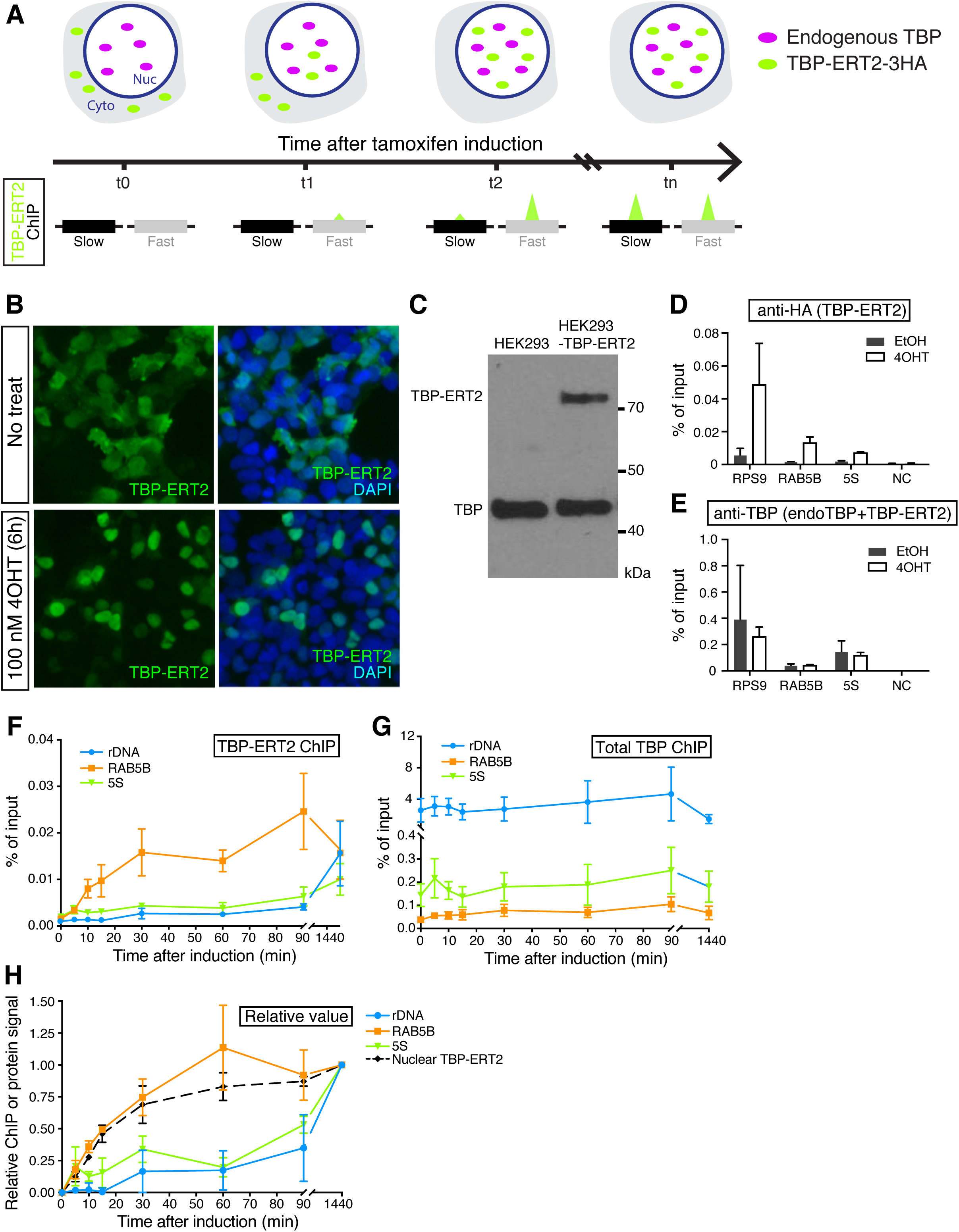
Nuclear translocation of TBP-ERT2. (*A*) Schematic illustration of tamoxifen-inducible, time-course ChIP analysis. (*B*) Western blot of whole cell extracts from HEK293 cells (parent) and TBP-ERT2 expressing cells (stable) with antibody against endogenous TBP. (*C*) Immunofluorescence image of TBP-ERT2 (green) and DAPI (blue) in HEK293 cells transiently transfected TBP-ERT2 expressing vector that are treated or non-treated with 100 nM tamoxifen (4OHT) for 6 hr. (*D*, *E*) Binding at the indicated loci (x-axis) of TBP-ERT2 (*D*) and total TBP (endogenous TBP and TBP-ERT2) (*E*) in cells stably expressing TBP-ERT2. Error bars indicate SEM (n=3). (*F, G*) Binding of TBP-ERT2 (*F*), total TBP (*G*) at the indicated loci at the indicated times after tamoxifen addition. (*H*) Abundance of nuclear TBP-ERT2 (black dashed line) and binding relative to the value at 1440 min. Error bars indicate SEM (n=3).

For measuring TBP binding dynamics in HEK293 cells, we expressed a HA_3_-tagged TBP derivative that contains ERT2, the ligand binding domain of the estrogen receptor that permits rapid nuclear translocation of the fused protein and binding to its target sites upon tamoxifen addition. As expected, tamoxifen induces translocation of TBP-ERT2 in the cytoplasm to the nucleus (Fig. 1B). We established a cell line that expresses TBP-ERT2 at a level that is 20% of endogenous TBP (Fig. 1C) and confirmed a significant increase of TBP-ERT2 association with target sites upon tamoxifen treatment (Fig. 1D). In contrast, target site association of total TBP (sum of endogenous TBP and TBP-ERT2) is similar before and after induction (Fig. 1E). The increase of the nuclear TBP-ERT2 amount starts within 5 minutes and is near steady-state within 30-60 minutes (Fig. 1F and Supplemental Fig. 1A, B), a time-frame comparable to that of *GAL1* induction in yeast.

### Different TBP turnover rates at Pol I, Pol II, and Pol III promoters

As an initial experiment, we analyzed promoters representing the three different RNA polymerases: rDNA (Pol I promoter); RAB5B (Pol II promoter); 5S (Pol III promoter). TBP-ERT2 association at the RAB5B promoter quickly increases in a manner kinetically indistinguishable from the increase in nuclear TBP-ERT2 concentration (Fig. 1H), indicating a turnover rate of < 5 minutes. In contrast, TBP-ERT2 association at the rDNA and 5S promoters increases much more slowly, indicating a much slower turnover rate at Pol I and Pol III promoters (Fig. 1F, H). The slow turnover of TBP-ERT2 at the rDNA promoter is not due to slow access to the nucleolus (the site of rDNA genes), because TBP-ERT2 localizes to the nucleolus at early time points when TBP-ERT2 is not bound at the rDNA promoter (Supplemental Fig. 1C). In contrast to TBP-ERT2, the ChIP signal for total TBP at all these promoters remains constant throughout the time course (Fig. 1G). The fast turnover at Pol II promoters and slow turnover at Pol I and Pol III promoters in human cells is similar to what is observed in yeast (van Werven et al. 2009; Zaidi et al. 2017).

### Genome-wide analysis of TBP binding dynamics

We performed time-course ChIP-seq analysis of TBP-ERT2 to examine the TBP binding dynamics at target sites on a genome-wide scale. To account for differences in immunoprecipitation efficiency among the samples, we added a constant amount of sonicated yeast chromatin prepared from a strain expressing HA-TBP as a spike-in control for each sample (Bonhoure et al. 2014; Orlando et al. 2014). We obtained 30-60 million reads/sample (Supplementary Table 1) and analyzed the 13,148 TBP-TERT2 peaks that met our criteria (Methods). TBP-ERT2-HA ChIP signals are thus normalized with yeast HA-TBP signals at each time point. ∼85% of TBP-ERT2 peaks are located in promoter regions (−1 kb to +100 bp from TSS) of annotated genes (Supplementary Fig. 2A); we presume the remaining 15% of target sites are mis- or un-annotated promoters. As expected, levels of TBP-ERT2 association at all time points (except 0) are highly correlated, and they gradually increase over time (Fig. 2A and Supplementary Fig. 2B, C). Most peaks show >4-fold increased association between the final induced state compared to the non-induced state (Supplementary Fig. 2D).

**Figure 2.**
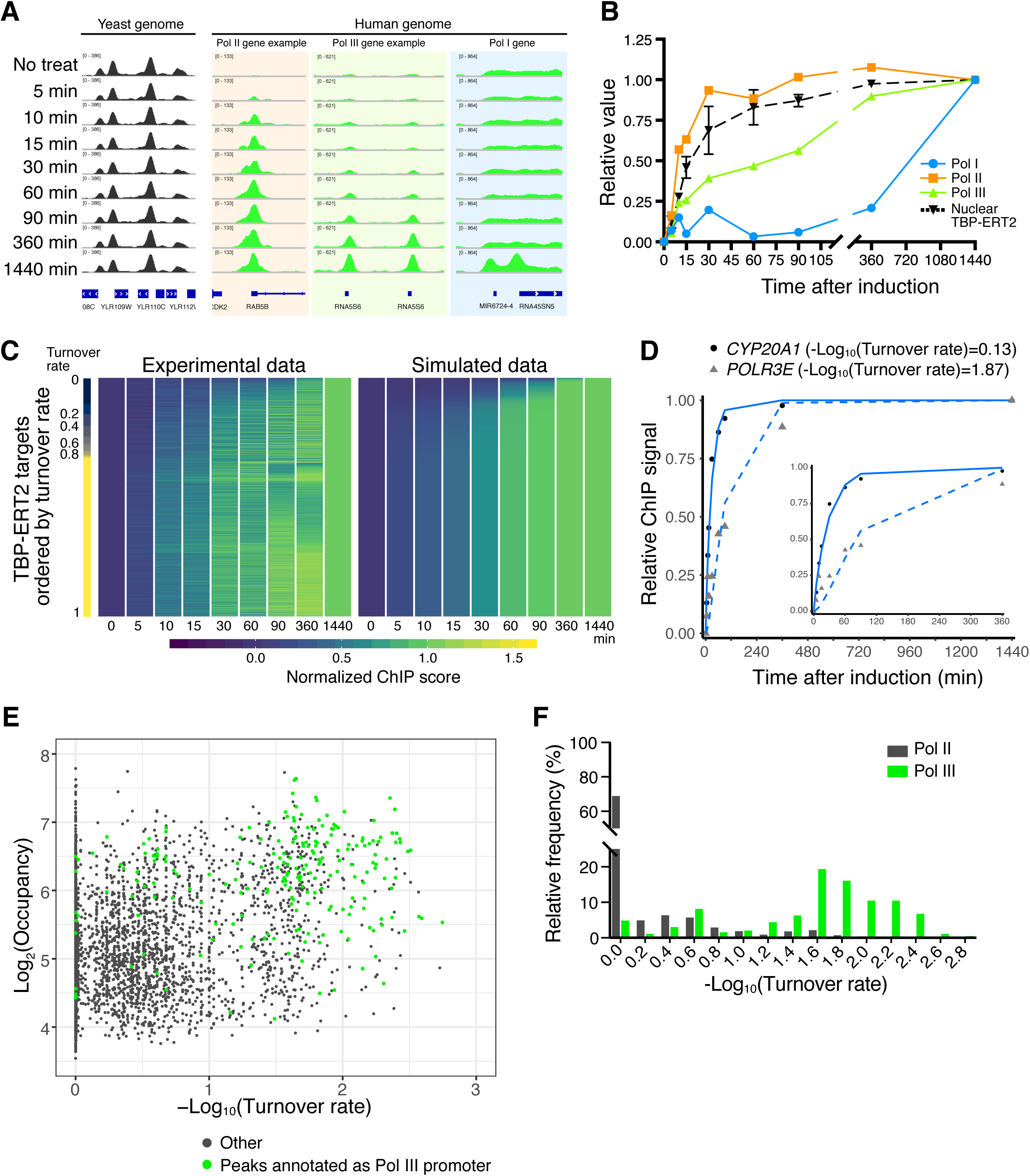
Time-course ChIP-seq analysis. (*A*) Examples of human loci bound by TBP-ERT2 and yeast loci bound by HA-TBP (spike-in). (*B*) TBP-ERT2 binding to the composite average of Pol I, Pol II, and Pol III promoters relative to the 1440 min sample. (*C*) TBP-ERT2 binding relative to the 1440 min sample (left) and simulated relative TBP-ERT2 binding by using turnover rate (right) of all detected peaks. Peaks are ordered by turnover rate value (Lambda). (*D*) Examples of TBP-ERT2 binding and simulated data at the CYP20A1 (solid line) and POLR3E (dashed line) loci. (*E*) Scatter plot of log2 TBP-ERT2 occupancy vs –log10 TBP-ERT2 binding turnover rate. (*F*) Histogram of the relative frequency of –log10 TBP-ERT2 binding turnover rate.

To compare the occupancy of TBP-ERT2 at each locus over time, we first subtracted ChIP signals at 0 min from those at all the other time points, and then calculated relative ChIP signals with respect to the endpoint (1440 min). On an overall basis and in accord with results on individual promoters, TBP-ERT2 occupancy at the majority of Pol II promoters is kinetically indistinguishable from nuclear translocation, whereas occupancy at Pol III (half-maximal occupancy at 90 minutes) and Pol I (half-maximal occupancy > 6 hr) is much slower (Fig. 2B).

### TBP turnover rates among Pol II or Pol III promoters are not related to occupancy

Using relative nuclear TBP-ERT2 levels (Supplementary Fig. 3A) and time-course ChIP-seq data, we employed a mathematical model (described in Methods) to calculate binding turnover rates (*λ*) for each TBP-ERT2 target site. The different kinetics of TBP-ERT2 occupancy can be simulated by using different binding turnover rates (Supplementary Fig. 3B). Slower turnover rates (larger value of –log10(turnover rate)) indicates that a longer time is required for the ChIP signal to reach to half-maximal steady-state level (Supplementary Fig. 3C). We extracted 6,476 sites for which the goodness of fit (squared norm of the residual) is < 0.4 and used this subset for subsequent analysis (Supplementary Fig. 4). Overall, the experimental data is captured well by the simulated data (Fig. 2C, D). Although the 5S and rDNA genes are highly repetitive with individual copies being heterogeneous in transcriptional activity and chromatin state (Conconi et al. 1989; Cloix et al. 2002), the time-course data at these genes fits the mathematical model arguing that the results are limited to active genes and against the idea of sub-populations of loci with different TBP binding dynamics.

Importantly, TBP-ERT2 binding turnover rates among Pol II promoters or among Pol III promoters can vary considerably, and these different turnover rates correlate poorly (r = 0.32) with the level of TBP-ERT2 occupancy (Fig. 2E, F). Thus, the TBP turnover rate at human promoters is an independent parameter that cannot be determined from the occupancy at one time point (Fig. 2E and Supplementary Fig. 3D).

### Slow TBP turnover rates at Pol II promoters are linked to a consensus TATA sequence, certain transcription factors, and accessible chromatin

To address why some Pol II promoters have slower TBP binding turnover rates than others, we categorized TBP-ERT2-bound Pol II promoters into three classes based on their turnover rates (Fig. 3A), and searched for enriched DNA sequence motifs that differentiated the classes. Interestingly, promoters with slow TBP turnover rates are strongly enriched for the TBP consensus binding motif namely the TATA element (Fig. 3B). To independently verify this observation, we scanned each peak region for the sequence with the highest match score to the position weight matrix of the TBP consensus motif. The cumulative curve and the box plot of match scores indicates that the slow turnover sites have higher similarity to the TATA motif, the middle turnover sites have intermediate similarity, and the fast turnover sites have lowest similarity (Fig. 3C, D). Half of the slow turnover sites have at least one strong TBP consensus motif in the same direction of the downstream gene (forward motifs; Fig. 3E). Forward motifs in the middle or slow turnover classes are densely clustered around ±100 bp around the transcription start site (TSS) of downstream genes, but forward motifs in the fast class or reverse motifs in all classes are more dispersed (Fig. 3F). In contrast to the TATA motif, other core promoter elements including the initiator are not enriched among any of the various turnover classes (Supplementary Fig. 5).

**Figure 3.**
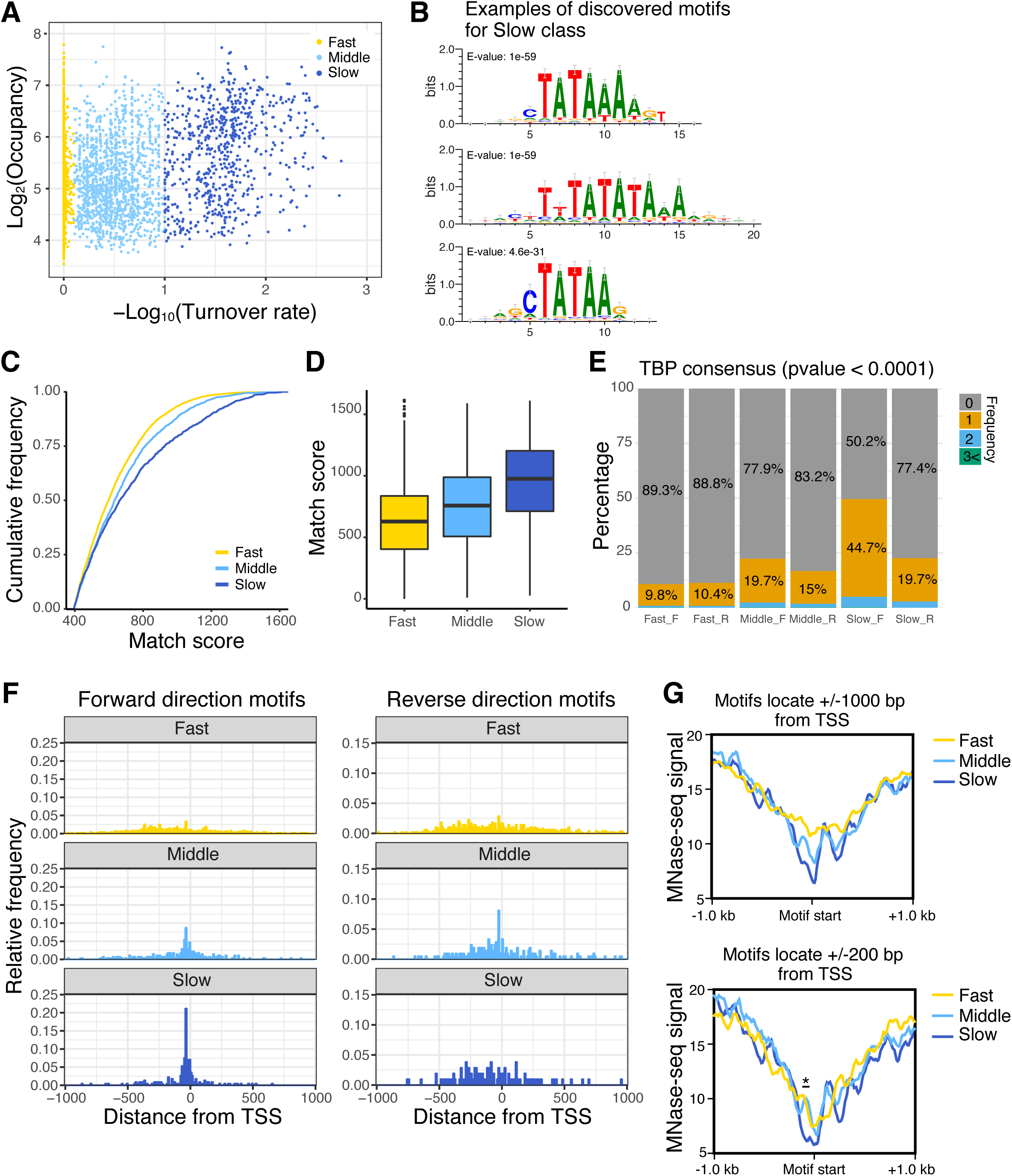
Pol II promoters with slow TBP turnover preferentially contain the consensus TATA sequence and fewer nucleosomes. (*A*) Scatter plot of TBP-ERT2 peaks annotated as Pol II promoters in log2 TBP-ERT2 occupancy vs –log10 TBP-ERT2 binding turnover rate. Colors indicate the classes categorized based on the binding turnover rates. (*B*) Top 3 DNA motifs enriched in the slow turnover class. (*C*) Cumulative frequency of match score to the position weight matrix of TATA consensus motif. (*D*) Boxplot of TATA consensus match scores to a forward-oriented consensus sequence for the indicated classes of promoters. (*E*) Percentage of promoters that have strong (*p* < 0.0001) hit sequences to TATA consensus motif, with colors indicating the number of TATA sequences in each promoter in the forward (F) or reverse (R) direction with respect to the downstream genes. (*F*) Position of TATA sequences in forward or reverse direction within 1 kb upstream or downstream of the TSS. (*G*) MNase-seq signal around hit sequences located within 1 kb upstream or downstream of the TSS.

To identify DNA-binding transcription factors that are linked to promoters with slow TBP turnover, we examined promoter regions (1 kb region upstream of the TSS) for presence of all known TF binding motifs (JASPAR database). Specifically, we searched for motifs that are enriched in the Slow class promoters as compared to those in Fast class promoters. Interestingly, motifs of SP1, KLF family members, NRF1, and NFYB are ranked in the top 10 based on the significance (E-values, Supplementary Fig. 6).

Nucleosome occupancy determined by MNase-seq ±1000 or ±200 bp from the TBP consensus motifs (MNase-seq data) is lowest in the slow turnover class and highest in the fast turnover class (Fig. 3G). As TBP can bind in a transcriptionally productive fashion to both TATA-containing and TATA-less promoters (which represent respectively 24% and 76% of human promoters (Yang et al. 2007)), these results indicate that TATA-containing promoters with low nucleosome occupancy are enriched in the slow turnover class. In addition, slow-class Pol II promoters that lack a strong TATA consensus motif tend to have higher AT content in the region just upstream of the TSS compared to other classes (Supplementary Fig. 7).

### High transcriptional activity is linked to both high TBP occupancy and slow TBP turnover

To investigate the relationships between TBP binding turnover rate, TBP occupancy, and transcriptional activity, we further subdivided TBP peaks at Pol II promoters into 6 groups based on TBP binding turnover rate (fast, middle, and slow) and occupancy (high and low) (Fig. 4A-C; for simplicity, analysis was restricted to 3,326 promoters having a sole annotation). Although the correlation between occupancy and turnover rate is weak, average TBP occupancy is slightly higher in the slow class (Fig. 4B). Occupancy of TAF1 is strongly correlated to TBP occupancy among the three classes (Fig. 4B, C), presumably because it an obligate component of the TFIID complex and hence behaves similarly to TBP. Strikingly, promoters with high TBP occupancy and slow TBP turnover have much higher Pol II occupancy than observed in the other five classes (Fig. 4D). Increased Pol II occupancy occurs regardless of the CTD phosphorylation state, suggesting that each stage of transcription is enhanced in this group. Importantly, Pol II occupancy at the other five classes is fairly similar, although it is slightly higher as a function of TBP occupancy and turnover rate.

**Figure 4.**
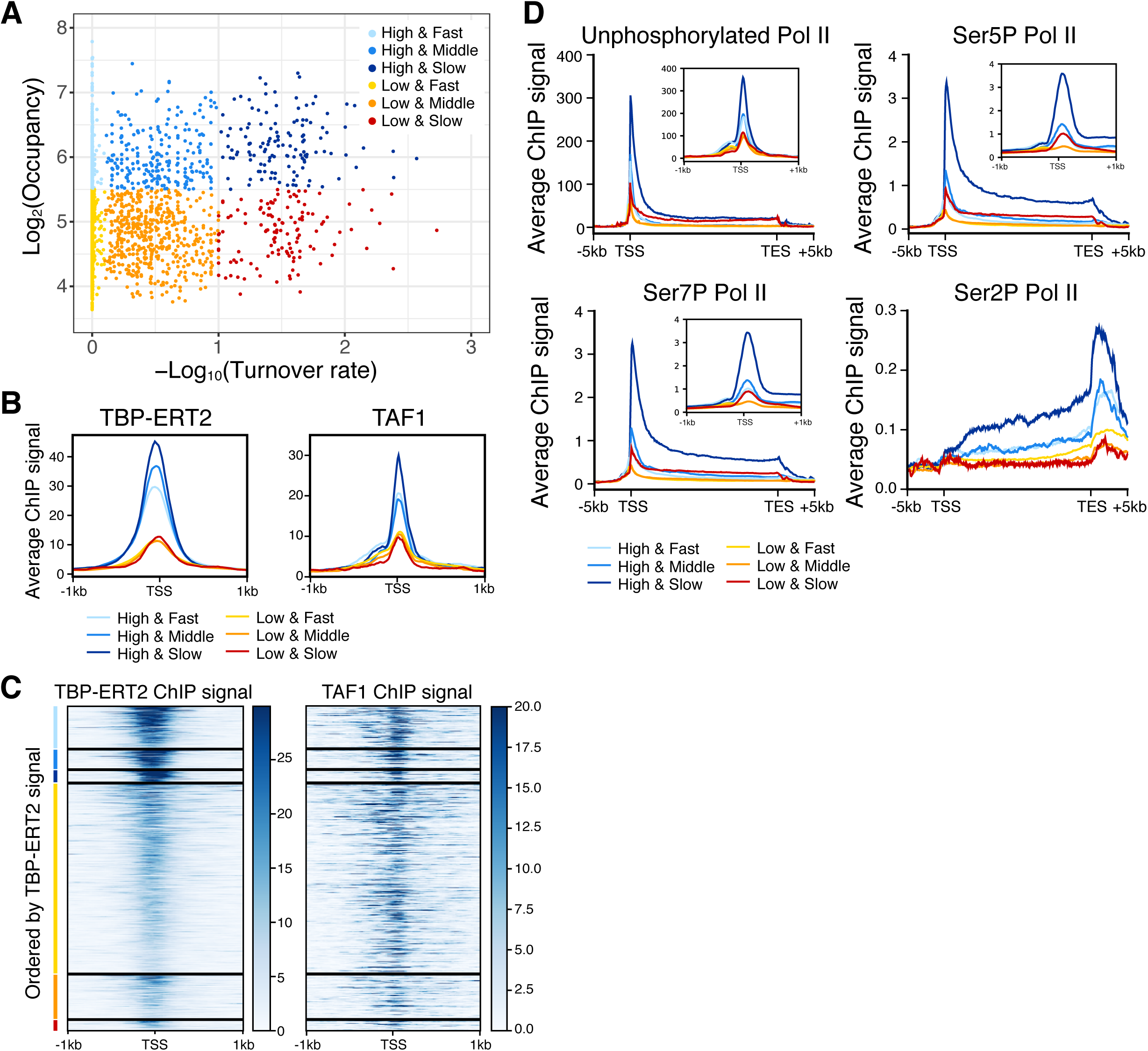
Strong transcription activity of genes with promoters with slow TBP turnover. Colors indicate the groups categorized based on the binding turnover rates and occupancy (n=3,326, High and Fast n=441, High and Middle n=202, High and Slow n=126, Low and Fast n= 1,985, Low and Middle n=462, Low and Slow n=110). (*A*) Scatter plot of TBP-ERT2 peaks annotated as Pol II promoters in log2 TBP-ERT2 occupancy vs –log10 TBP-ERT2 binding turnover rate. Colors indicate the groups based on the binding turnover rates and occupancy. (*B*) Heatmap of TBP-ERT2 binding at each target site within the indicated groups. (*C*) Means of TBP-ERT2 binding between −1 kb to +1 kb from the TSS in each group. (*D*) Means of unphosphorylated or phosphorylated Pol II binding between the TSS and TES plus 5 kb upstream or downstream in each group. Plots in the small windows are Means of TBP-ERT2 ChIP-seq signal along −1 kb to +1 kb from TSS in each group. (*E*) TBP-ERT2 and TAF1 binding to Pol II promoters (ordered by TBP-ERT2 ChIP signal) between −1 kb to +1 kb from the TSS.

The high transcriptional activity of promoters with high TBP occupancy and slow TBP turnover is confirmed by PRO-seq experiments (Woo et al. 2018) that measure the amount of nascent RNA transcripts (Supplementary Fig. 8A-C). All six classes have a similar Pol II pausing index that represents the ratio of paused Pol II in the promoter-proximal region relative to the elongating Pol II throughout the coding region (Supplementary Fig. 8D). In addition, the high TBP occupancy and slow TBP turnover group has a low antisense:sense ratio compared to the other classes, indicating a strong preference for transcription in the sense direction in this group (Supplementary Fig. 8E, F). Thus, transcriptional activity is not simply dependent on TBP occupancy, but rather is also strongly associated with TBP binding dynamics.

### TBP binding dynamics is differentially associated with recruitment of chromatin-modifying factors

Using the same six classes of promoters, we analyzed the relationship of TBP occupancy and dynamics with respect to the recruitment of chromatin regulatory factors. Interestingly, GCN5(KAT2A) histone acetylase (catalytic subunit of SAGA and other complexes) and BRG1 (catalytic subunit of SWI/SNF and related nucleosome-remodeling complexes) are specifically enriched at promoters with slow TBP binding turnover (Fig. 5A). This result is observed at both high and low TBP occupancy, although recruitment of these chromatin-modifying complexes is lower at promoters with low TBP occupancy. In contrast, recruitment of p300, a histone acetylase, does not vary as a function of TBP dynamics (Fig. 5A).

**Figure 5.**
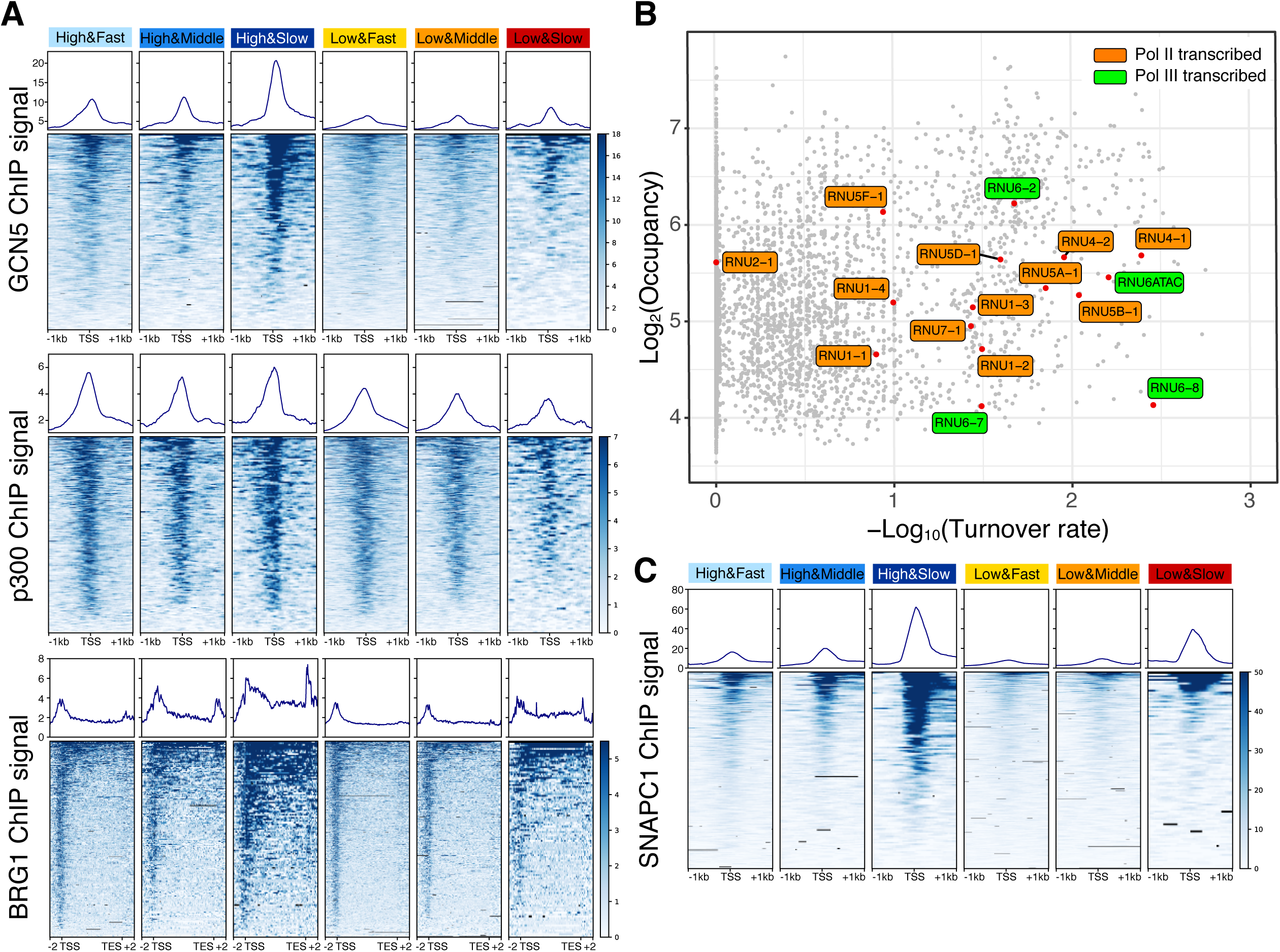
GCN5, BRG1 and SNAPC1 are enriched in genes with slow TBP binding dynamics. (*A*) Means and heatmaps of GCN5/KAT2A (top), p300 (middle), and BRG1 (bottom) binding on each target site. (*B*) Distribution of snRNA genes (red) in the scatter plot of log2 TBP-ERT2 occupancy vs –log10 TBP-ERT2 binding turnover rate. Gray and blue respectively indicates Pol II and Pol III transcribed snRNA. (*C*) Means and heatmaps of SNAPC1 binding on each target site.

### snRNA promoters recognized by the SNAPc complex have slow TBP turnover

snRNA genes are transcribed by either Pol II or Pol III. All snRNA promoters contain a proximal sequence element (PSE) recognized by the snRNA-activating protein complex (SNAPc) that consists of five subunits. Consistent with typical Pol III-transcribed genes, slow TBP binding turnover is observed at the Pol III-transcribed snRNA genes RNU6 and RNU6ATAC. In addition, Pol II-transcribed snRNA genes (except for RNU2) also show slow TBP binding turnover (Fig. 5B). These results suggested that slow TBP binding turnover at snRNA genes may be related to SNAPc recruitment. To test this hypothesis, we examined the six classes of promoters for binding of SNAPC1, a component of SNAPc, that binds to highly active Pol II genes in addition to snRNA genes. SNAPC1 is strongly enriched at promoters with slow TBP turnover (Fig. 5C), strongly suggesting that SNAPc contributes to slow TBP dynamics at both Pol II and Pol II promoters.

### Differential TBP turnover at Pol III promoters

Although TBP binding turnover on Pol III promoters is generally slower than on Pol II promoters, several Pol III promoters show fast turnover at speeds comparable to that of typical Pol II promoters. Almost all Pol III promoters that show fast TBP binding turnover are tRNA promoters, except for one case annotated as a U6 RNA pseudogene (Fig. 6A). To address the basis of differential TBP turnover at Pol III promoters, we sub-divided all Pol III promoters into three classes based on turnover rate and analyzed TBP occupancy and chromatin structure (MNase-seq and DNase-seq). Pol III promoters with slow TBP turnover have more open chromatin structure (higher DNase-seq signal and lower MNase-seq signal; Fig. 6B, C) as compared to Pol III promoters with faster TBP turnover. In this regard, Pol III promoters behave similarly to Pol II promoters.

**Figure 6.**
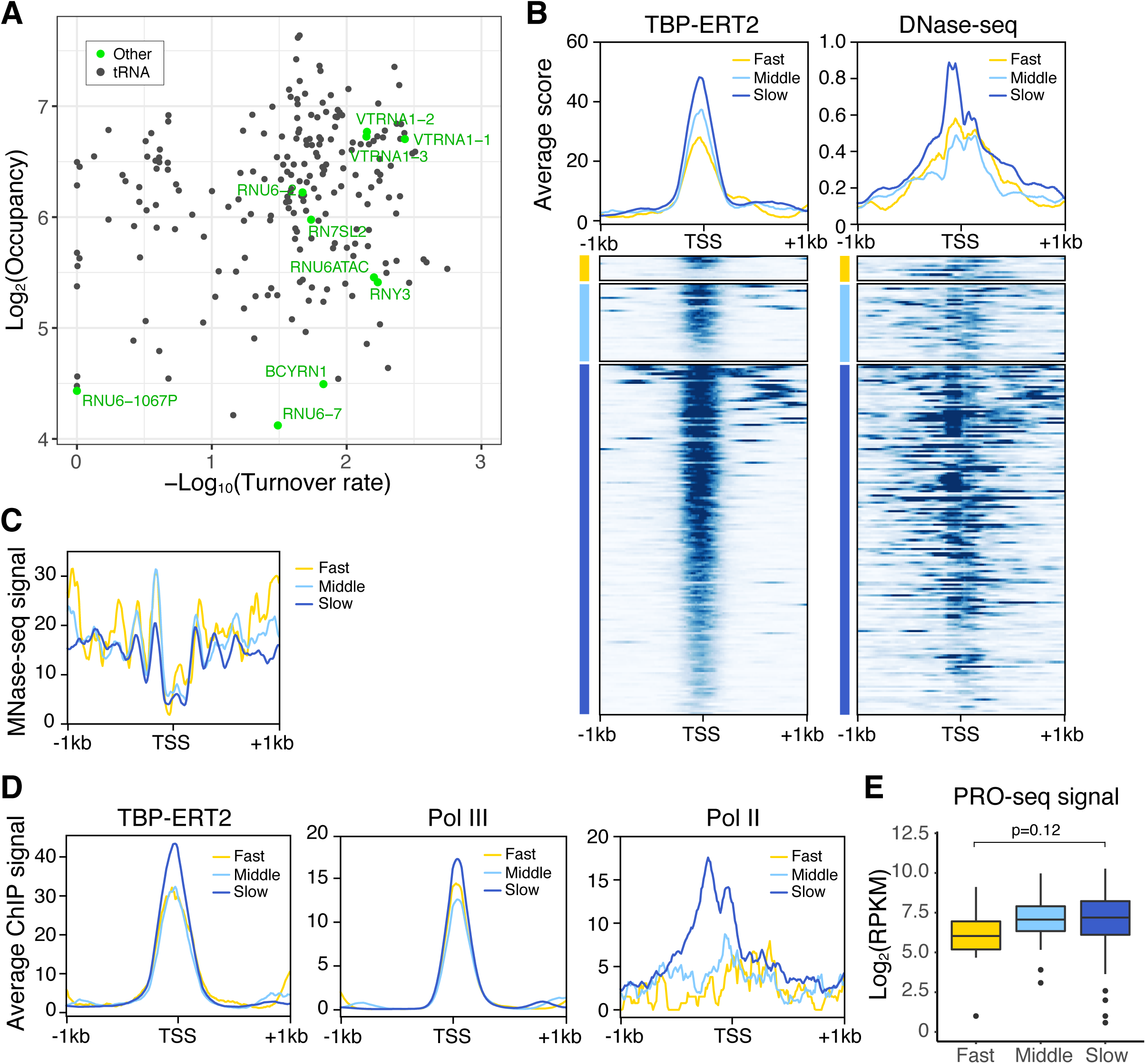
Pol III promoters with slow TBP binding dynamics have less nucleosomes and higher Pol III recruitment. (*A*) Scatter plot of TBP-ERT2 peaks annotated as Pol III genes in log2 TBP-ERT2 occupancy vs –log10 TBP-ERT2 binding turnover rate. (*B*) Means and heatmaps of TBP-ERT2 binding DNase-seq signal on each target site near Pol III genes. (*C*) Means of MNase-seq signal along −1 kb to +1 kb from the TSS in each class. (*D*) Means of Pol III (POLR3A) binding along −1 kb to +1 kb from the TSS in each class. (*E*) Boxplot of PRO-seq signal at Pol III genes in each class.

We investigated the association between the transcriptional activity and TBP occupancy or turnover rate with a subset of Pol III promoters (n=130; for simplicity, we restricted these to those having a sole annotation > 1 kb from other genes). In contrast to the case of Pol II promoters, Pol III occupancy and RNA levels are quite similar between the tRNA promoters with slow or fast TBP turnover (Fig. 6D, E), and the consensus TATA sequence occurs equivalently on Pol III promoters of both turnover classes (Supplementary Fig. 9). The subtle increased levels of Pol III on the slow class promoters is likely due to slight increases in the level of TBP occupancy. Although Pol II binds close to active, but not inactive, Pol III promoters in human cells (Moqtaderi et al. 2010; Oler and Cairns 2010; Raha et al. 2010), Pol II enrichment is unexpectedly higher in the slow class promoters than the other classes (Fig. 6D).

## DISCUSSION

### A ERT2-based, time-course ChIP method to measure dynamics of transcription factor binding at target sites on a genome-wide scale in human cells

Although transcription factor binding to chromatin is highly dynamic on an overall basis, binding dynamics at target loci in metazoan cells have not been investigated. Here, we describe an ERT2-based, inducible time-course ChIP-seq method to address the dynamics of transcription factor binding at target loci on a genome-wide scale. This method should be applicable to all transcription factors whose function is retained upon fusion to the ERT domain that mediates nuclear import upon treatment with tamoxifen. The turnover rate measured by this approach is determined by the dissociation rate of the endogenous transcription factor bound to promoters prior to tamoxifen addition and the association rate of the tamoxifen-induced ERT2 derivative. While our approach cannot directly distinguish between association and dissociation rates, we strongly suspect that dissociation rate is the primary determinant. In particular, slow turnover in all cases tested is associated with high chromatin accessibility, the opposite of what one would expect if association rates were important.

### Relative TBP turnover rates at Pol I, Pol II, and Pol III promoters are evolutionarily conserved

Although TBP is required for transcription by all three nuclear RNA polymerases, the transcription machineries used by each of these RNA polymerases are otherwise very different. In general, our results indicate that TBP turnover in human cells is very rapid at Pol II promoters, considerably slower at Pol III promoters, and extremely slow at Pol I promoters. This observation is remarkably similar to what occurs in yeast cells (van Werven et al. 2009), indicating that relative TBP turnover rates at promoters transcribed by the three different transcriptional machineries is conserved from yeast to human. Although the Pol II and Pol III machineries and the relevant TBP complexes are highly conserved across eukaryotes, the yeast and human Pol I-specific TBP complexes are very different. Aside from TBP, the proteins in the human SL1 complex (Comai et al. 1992) and have only weak sequence similarities with those in the yeast Core factor (Lin et al. 1996) even though that both complexes recognize the core promoter element in the rDNA. The exceptionally low turnover rates of such different TBP complexes at the rDNA promoter strongly suggests that low TBP turnover is fundamentally important at Pol I promoters. As rDNA genes exist in long tandem copies, only some of which are active at any one time, this suggests that the active and inactive states of rDNA promoters are maintained for considerable time.

Although the induction kinetics of nuclear TBP are similar in yeast and human (saturation by 30-60 minutes), induced human TBP requires a longer time than yeast TBP to saturate TBP occupancy levels on Pol I and Pol III promoters (Pol I: > 3 h in yeast and > 24 h in human; Pol III: 60-90 min in yeast and > 6 h in human) (van Werven et al. 2009). We suspect that this difference reflects the much longer time for cell division in yeast and human cells. Although the molecular basis for this difference is unclear, it might be due to clearance of DNA-bound TBP during DNA replication.

### Determinants of fast vs. slow TBP turnover rates among Pol II or Pol III promoters

Although Pol II promoters with slow TBP turnover tend to have higher TBP occupancy, TBP turnover rate at Pol II promoters is poorly correlated (r = 0.31) with TBP occupancy in human cells. At most Pol II promoters, the kinetics of TBP-ERT2 binding is indistinguishable from that of TBP-ERT2 translocation into the nucleus upon tamoxifen induction, indicating a very fast turnover rate. However, some Pol II promoters have a much slower turnover rate as well as distinct properties from the majority class of promoters with fast TBP turnover rates. Pol II promoters with slow TBP turnover 1) frequently contain consensus TATA motifs, 2) are associated with high levels of transcription of downstream genes, 3) have low nucleosome occupancy, and 4) show stronger recruitment of chromatin-modifying activities such as GCN5 histone acetylase and the SWI/SNF nucleosome remodeling complex. snRNA promoters, which specifically rely on SNAPc for transcription, also exhibit slow TBP turnover. These results demonstrate that TBP binding dynamics, not simply TBP occupancy, is important for transcriptional activity and chromatin structure around promoters.

These observations on TBP binding dynamics on Pol II promoters in human cells differ from what occurs in yeast cells. Yeast promoters with slow TBP turnover were reported to have higher mRNA levels (van Werven et al. 2009), but higher levels of Pol II occupancy on these genes is not evident in the same paper. Furthermore, a reanalysis of the same data using a different model and gene set suggested the opposite conclusion that yeast promoters with fast TBP turnover have modestly higher levels of nascent transcription (Zaidi et al. 2017). In addition, TATA-containing yeast promoters are not enriched in the slow TBP turnover class and have modestly faster binding turnover of TBP compared to TATA-less promoters (van Werven et al. 2009; Zaidi et al. 2017).

Both Pol II and Pol III promoters with relatively slow TBP turnover rates are associated with more open chromatin structure. However, unlike the case for Pol II promoters, there is little if any relationship between turnover rates and transcriptional activity on Pol III promoters. Interestingly, and in accord with the fact that Pol II binds closely to active, but not inactive, tRNA promoters in human cells (Moqtaderi et al. 2010; Oler and Cairns 2010; Raha et al. 2010), tRNA promoters with slow TBP turnover are associated with higher levels of Pol II binding. As tRNA and most other Pol III genes lack TATA sequences in their promoters, recruitment of Pol II and associated chromatin-modifying activities, might provide an alternative mechanism to ensure stable TBP binding.

### Implications for transcriptional mechanisms

TFIID binds to promoters via sequence-specific interactions between TBP and the TATA sequence as well as interactions of TAFs with initiator and downstream promoter elements. The simplest explanation for the properties of promoters with slow TBP turnover is that TBP interaction with a consensus TATA element stabilizes the TFIID complex on the promoter. As such, the weaker TBP-TATA interaction at so-called TATA-less promoters would lead to faster dissociation rates. In this regard, the transcription factors preferentially linked to Pol II promoters with slow TBP turnover might directly recruit TFIID to promoters and hence stabilize the TFIID-TATA interaction. Although Mediator is the direct target of most activators, there is a subset of activators in yeast (Kuras et al. 2000; Li et al. 2000) and human (Chen et al. 2013) cells that directly recruit TFIID.

A strong TBP-TATA interaction is important for high levels of transcription mediated by most activator proteins (Iyer and Struhl 1995; Lee and Struhl 1995; Struhl 1996), presumably because it is associated with increased transcriptional reinitiation (multiple rounds of transcription following preinitiation complex formation) (Yean and Gralla 1997; Yean and Gralla 1999; Yudkovsky et al. 2000; Wong et al. 2014) and transcriptional bursting (Zenklusen et al. 2008; Sanchez and Golding 2013). Gene looping between promoter and terminator regions is likely to facilitate pol II recycling after transcription termination (O’Sullivan et al. 2004; Tan-Wong et al. 2012), and this may contribute to increased transcriptional reinitiation at promoters with slow TBP turnover. In this regard, we observed increased Swi/Snf occupancy at the 3’ regions of genes whose promoters show slow TBP turnover, similar to Swi/Snf occupancy at 3’ regions of genes observed in *Arabidopsis* (Archacki et al. 2017). The function of transcriptional activator proteins is linked to reduced nucleosome occupancy via recruitment of chromatin-modifying activities, and this might account for why promoters with slow TBP turnover show decreased nucleosome occupancy. Thus, slow TBP turnover results in increased transcription beyond the contribution of TBP occupancy.

The differences between yeast and human Pol II promoters with respect to TBP dynamics are likely to reflect differences between transcriptional mechanisms in these two organisms. First, at human (and most other eukaryotic) promoters, Pol II pauses just downstream of the initiation site, but this pause does not occur in yeast cells because it lacks NELF, a critical factor for the pause (Adelman and Lis 2012; Zhou et al. 2012; Kwak and Lis 2013). This Pol II pause helps keep the chromatin open in the vicinity near the pause (Gilchrist et al. 2010). Second, yeast TBP exists in two distinct active forms for Pol II transcription, TFIID and a form lacking TAFs (probably free TBP) (Kuras et al. 2000; Li et al. 2000), whereas human TBP is predominantly (and probably exclusively) in the form of TFIID. One possible explanation is that yeast TBP has a faster turnover at yeast Pol II promoters than TFIID, thereby accounting for why TATA-containing promoters (which are favored by the free TBP form) have faster TBP turnover than TATA-lacking promoters. In human cells, where TFIID is presumably the exclusive form for TBP, so slow turnover is simply due to increased binding affinity via the TBP-TATA interaction.

Our results showing that higher transcriptional activity is linked to slow TBP turnover beyond simple TBP occupancy are consistent to results with yeast Rap1 (Lickwar et al. 2012). We suggest that relationship reflects the fact that assembly of active transcription complexes takes some time. Proteins with fast turnover might initiate the transcription process, but will be unable to complete it if they dissociate prior to full assembly of an active complex. Thus, while factor occupancy is clearly linked to transcriptional activity, the amount of time that the factor remains associated with the chromatin template has an independent effect that could be manifest on the preinitiation complex *per se* or transcriptional reinitiation that results in multiple transcripts for each assembled complex.

## MATERIALS AND METHODS

### Plasmid construction

The DNA fragment coding 3xHA was prepared by annealing oligos (Supplementary Table 3,) and inserted between the *Bam*HI and *Eco*RI sites of pBudCE4.1 (Invitrogen). The ERT2 DNA fragment was PCR amplified from pCAG-Cre-ERT2 (Addgene #14797) using 1Fw and 1Rv primers and inserted between *Sal*I and *Bam*HI sites of pBudCE4.1-3xHA. ERT2-3HA fragment was then PCR amplified using 2Fw and 2Rv primers and inserted between *Bmt*I and *Bam*HI sites of pmCherry-C1 (Clontech). The TBP coding sequence was PCR amplified from human cDNA and was cloned into *Sal*I site of this plasmid containing ERT2-3HA, yielding pCMV-TBP-ERT2-3HA.

### Cell culture

pCMV-TBP-ERT2-3HA construct was transfected to HEK293 (ATCC) cells, and the stable line was established by G418 selection. Cells were routinely cultured in phenol-red free DMEM containing 10% FBS.

### Nuclear and nucleolar fractionation

Cells cultured in 6 cm dish were washed twice with ice-cold PBS twice, then scraped and collected in 1.5 ml tubes. After brief centrifugation, cells were suspended in 250 μl of CE buffer (10 mM Hepes-KOH (pH7.5), 1.5 mM MgCl2, 10 mM KCl, 0.05% NP-40) and incubated for 3 min on ice. Cells were again centrifuged for 3 min at 3,000 rpm, and the pellet were suspended in 250 μl of RIPA buffer (50 mM Tris-HCl (pH8.0), 150 mM NaCl, 5 mM EDTA, 1% Triton X-100, 0.5% Na-Deoxycholate, 0.1% SDS) and sonicated (Misonix 3000, level 2, on 30 sec off 30 sec, total 2 min). After centrifugation for 12,000 rpm for 10 min, the supernatant was collected and used as the nuclear fraction.

To isolate nucleoli, cells in five 150-mm dishes were trypsinized and collected in the 15 ml tubes, then suspended in ice-cold CE buffer without detergent (10 mM Hepes-KOH (pH7.9), 1.5 mM MgCl_2_, 10 mM KCl, 0.5 mM DTT) and incubated for 5 min on ice. The cell suspension was transferred to pre-chilled 7 ml Dounce tissue homogenizer, and homogenized to obtain nuclei. After centrifuge and removed supernatant, the nuclear pellet was suspended in 3 ml of S1 buffer (0.25 M Sucrose, 10 mM MgCl_2_) and then put on the top of 3 ml S2 buffer (0.35 M Sucrose, 0.5 mM MgCl2). The tube was centrifuged at 2,500 rpm for 5 min at 4°C, and after that supernatant was removed. The pellet was suspended in 3 ml of S2 buffer and sonicated for 3 x 10 sec using a probe sonicator. The sonicated solution was layered onto 3 ml of S3 buffer (0.88 M Sucrose, 0.5 mM MgCl_2_), and centrifuged for 10 min at 3,500 rpm, 4°C. The pellet containing nucleolus was washed with 500 μl of S2 buffer, spun down at 3,500 rpm for 5 min and then used for the assay.

### Sample preparation during the time course

TBP-ERT2 nuclear translocation was induced by 4OHT (Sigma, H7904) addition at the final concentration of 100 nM to culture medium. Cells were retained in the CO2 incubator for indicated times. For western blotting, cells were immediately put on ice, and proceed to the subcellular fractionation. For ChIP, culture medium was removed and then exchanged by fixing solution (see below) at room temperature to avoid over crosslinking. This step takes 2.5 minutes, thus we summed up this time gap and incubation time and used as real time points (7.5, 12.5, 17.5, 32.5, 62.5, 92.5, 362.5, 1442.5 min) for simulating turnover rate.

### Chromatin immunoprecipitation

Chromatin was prepared from two 150 mm dishes. After removing the culture medium, cells were fixed at room temperature with fixing solution (1% formaldehyde, methanol-free in DMEM without serum) for 10 min. Fixation was stopped by adding 125 mM glycine. Cells were washed with ice-cold PBS twice, and then scraped from dishes and collected. Cells were suspended in 2 ml Lysis Buffer 1 (50 mM Hepes-KOH (pH7.5), 140 mM NaCl, 1mM EDTA, 10% Glycerol, 0.5% NP-40, 0.25% Triton X-100, with protease inhibitor and phosphatase inhibitor), rotated for 10 min at 4°C. After collecting cells by centrifugation, cells were re-suspended in 2 ml Lysis Buffer 2 (10 mM Tris-HCl (pH8.0), 200 mM NaCl, 1 mM EDTA, 0.5 mM EGTA, with protease inhibitor and phosphatase inhibitor) and rotated for 10 min at 4°C. Cells were collected by centrifugation and re-suspended in 1.3 ml Lysis Buffer 3 (10 mM Tris-HCl (pH8.0), 100 mM NaCl, 1 mM EDTA, 0.5 mM EGTA, 0.1% Sodium deoxycholate, 0.5% N-Lauroylsarcosine, with protease inhibitor and phosphatase inhibitor), and then sonicated (Misonix 3000 sonicator, level 2, ON 30 sec, OFF 30 sec, 7 cycles (total sonication time is 3.5 min)). Majority of the fragment sizes were approximately 200-500 bp. Cell lysates were centrifuged at 12,000 rpm for 10 min at 4°C after adding Triton X-100 (1% final concentration), and the supernatants were used for the analysis. The protein concentration of the lysates was measured by BCA Protein Assay Kit (Thermo Scientific) and the amounts of the lysates yielding 1.3 µg total protein were used for immunoprecipitation. The volume of the lysates were adjusted to 1 ml by adding Lysis Buffer 3. For sequencing samples, yeast chromatin prepared from HA-TBP expressing strain were added at the constant ratio (0.039 µg total protein in the lysate, that corresponds to 3% of total human protein amount in the lysates) to the human cell lysates as a spike-in control. 1/10 volumes of the mixed lysates were kept as input samples. 0.3 µg of anti-HA antibody (Santa Cruz #sc-7392) or 3 µg of anti-TBP antibody (Abcam #ab51841) was added to the lysates, and the mixtures were rotated overnight at 4°C. Immunoprecipitation was performed at 4°C for 3 hr with 50 µl protein G dynabeads (Thermo Fisher). Beads were washed five times with ice-cold Wash buffer 1 (50 mM Hepes-KOH (pH7.6), 500 mM LiCl, 1 mM EDTA, 1% NP-40, 0.7% Sodium deoxycholate) and rinsed with TE (10 mM Tris-HCl (pH8.0), 1 mM EDTA) with 50 mM NaCl. Immunoprecipitates were eluted by incubating the beads in 75 µl Elution buffer (50 mM Tris-HCl (pH 8.0), 10 mM EDTA, 1% SDS) at 65°C for 15 min and collecting the supernatants. This step was repeated twice. The eluted samples were incubated overnight at 65°C for de-crosslinking. After treating with RNase A and Proteinase K, DNA was purified with PCR purification kit (QIAGEN) by adding 100 µl water. 55.5 µl of eluted DNA from IP samples and 2.5 µl of eluted DNA (diluted to 1/10) from input samples were used for the library preparation. For qPCR analysis, 3.3 µl of eluted DNA from IP samples and four dilution series of input samples for standards were used. Primers are listed in Table S3.

### Yeast chromatin preparation

HA-TBP yeast strain was grown in YPD at 30 °C to an 0.4 OD600 and fixed for 20 min at room temperature with final 1% formaldehyde. Then the fixation was quenched with final 250 mM glycine for 5 min at room temperature. Cells were collected and washed with PBS once, and were disrupted in Mini Bead Beater (BioSpec Products, Inc.) with 3 cycles of 5 min run at maximum speed and 1 min rest on ice following suspending in 400 µl of FA lysis buffer (50 mM Hepes-KOH (pH7.5), 150 mM NaCl, 1% Triton X-100, 0.1% Sodium deoxycholate, 0.1% SDS). After centrifugation and removal of the supernatant, the pellet was suspended in 480 µl of FA lysis buffer and sonicated (Misonix 3000 sonicator, level 6, ON 10 sec, OFF 10 sec, 21 cycles (total sonication time is 3.5 min) × 6 times with 1 min interval). The sample was centrifuged at maximum speed at 4°C for 20 min, and then the supernatant was collected and used as yeast chromatin.

### Library preparation and sequencing

Sequencing library preparation was performed with NEBNext Ultra DNA library prep kit (NEB #E7370S) and NEBNext Multiplex Oligos (NEB #E7335S) following the manufacturer’s instructions. PCR cycles for amplification of adaptor-ligated DNA were 9 cycles for input samples and 14 cycles for IP samples. Large fragments were removed using 0.4 × AMPure beads (Beckman Coulter). Libraries were sequenced on NextSeq 500.

### ChIP-seq computational analysis

We concatenated the genome sequences of human (hg38) and cerevisiae (sacCer3) to generate a combined genome sequence. We added “_s” to all yeast chromosome names to avoid chromosome name duplication, and built a custom Bowtie2 index from this combined genome sequence. Sequenced reads were aligned to this custom library with default parameters (--sensitive). The resulting SAM files were then split into the two files containing reads mapped to human chromosomes and mapped to yeast chromosome. Duplicates were removed by Picard, and only reads exceed mapping quality 30 were retained except for specific analysis (repetitive genes including ribosomal DNA). The number of reads mapped to human or yeast genomes and that passed the quality filter are listed in Supplementary Table 1. We calculated RRPM (reference adjusted RPM) by dividing the number of the reads mapped to human genome loci by the total reads number mapped to yeast genome (the numbers in the MQ>30 column in Supplementary Table 1, and that in the MQ>0 column for repetitive gene analysis). This RRPM value was used as the intensity of the ChIP signal in this study. Peak calling was performed using MACS2 with a p-value threshold of 0.01. We extracted 13,148 peaks that were called at least two samples in the later time points (60, 90, 360, 1440 min). Peak annotation was performed using UROPA with custom GTF file combined GRCh38.94 and tRNA GTF files downloaded from Ensembl and UCSC genome browser respectively.

### Modeling of protein translocation to the nucleus

We assume that the increase of nuclear fraction of TBP-ERT2 corresponds to decrease of cytoplasmic fraction of TBP-ERT2 at given time point under the condition with excess amount of 4OHT compared to the number of TBP-ERT2 molecules, and that the amount of total TBP-ERT2 does not change over time. This implies the following constraint

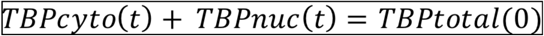

The dynamics of TBP can be described as follows

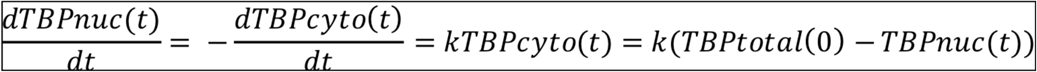

Thus,

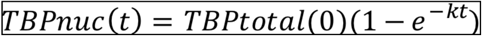

We consider the situation where all the TBP-ERT2 are initially in the cytoplasm, and eventually end up in the nucleus. Indeed,

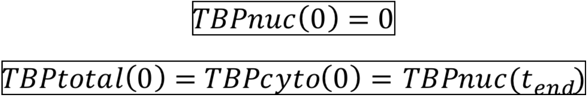

Using equation (5), equation (3) can be rewritten as:

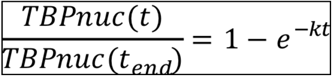

We assume that the measured nuclear TBP-ERT2 signal (mTBPnuc(t)) is sum of the true signal (TBPnuc(t)) and background (BG) derived from experimental errors, like imperfect separation of cytoplasmic and nucleus. Thus,

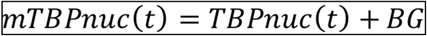

From equation (3),

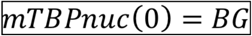

Indeed, equation (5) can be transformed as:

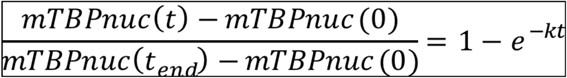

We measured the nuclear amount of TBP-ERT2 at all time points (0, 5, 10, 15, 30, 60, 90, 360, 1440 min) using western blotting analysis, and determined k in equation (8) that yields the best fit to the measured data using lsqcurvefit function of Matlab (Supplementary Fig. 5A.

### Turnover model

We used a simple model to describe the interaction between TBP and DNA with assumption that (1) TBP detected by western blotting is mostly free in the soluble fraction (2) turnover rate of endogenous TBP and TBP-ERT2 are same (3) when TBP molecule dissociates from DNA, the other TBP molecule immediately comes to occupy the binding site. Thus, the dynamics of DNA bound TBP-ERT2 can be described with TBP turnover rate *λ* as follow:

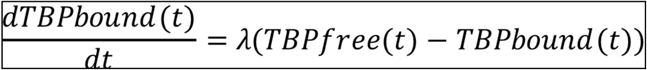

We also assume that the measured TBP-ERT2 ChIP signal (mTBPbound(t)) is sum of the true signal (TBPbound(t)) and background (BG). Thus,

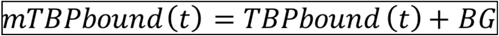

From equation (3), theoretically

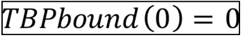

and thus,

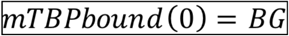

We define C(t) and W(t) as relative concentrations to endpoint of DNA bound TBP (measured by ChIP-seq) and TBP protein (Signal detected by western blotting) respectively.

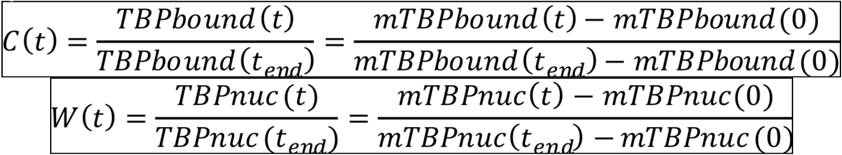

Indeed, equation(x) can be written with relative concentrations

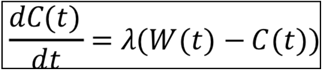

*λ* for each locus was determined using ODE45 and lsqcurvefit function of Matlab to yield the best fit to the measured data. We used multiple initiation points to avoid local minima. We excluded the peaks that show low goodness-of-fit (residual sum of squares > 0.4), and used the rest of the 6,476 peaks were used for the later analysis. We defined turnover rate as -log10(*λ*). In this model, the speed of the protein translocation to the nucleus is the major limitation for measuring TBP binding kinetics. As shown in the simulated result (Supplementary Fig. 3B), in the cases that TBP binding turnover rate exceeds 1, it is out of the range of the precise detection in our experimental system. Thus, we set a maximum turnover rate as 1 to lump together all the TBP target sites of which turnover rates exceed the detection limit as “very fast” category.

### Motif analysis

6,258 peaks which are not annotated as pol III promoters or genes were categorized into three classes based on their turnover rate: Fast 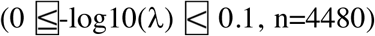, Middle 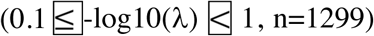, Slow 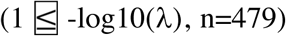. 217 peaks which are annotated as pol III promoters or genes were also categorized into three classes based on the same criteria: Fast (n=11), Middle (n=35), Slow (n=171). FASTA files of the DNA sequences of peaks in Fast class and Slow class were used as input for RSAT peak-motifs (http://rsat.sb-roscoff.fr/peak-motifs_form.cgi) to search Slow class enriched motifs. The match scores, p-values and locations of TBP consensus motifs (JASPAR MA0108.2), INR (JASPAR POL002.1 or Vo ngoc et al(Vo Ngoc et al. 2017)), BREu (JASPAR POL006.1), and BREd (JASPAR POL007.1) in the peaks were searched by PWMscan of PWMtools (https://ccg.epfl.ch/pwmtools/pwmscan.php) with scanning options (Score cut-off: 0). For the search of known TF binding motifs enriched in Slow class, −1-kb region from TSS of genes classified as Fast and Slow class were used as input for CentriMo (http://meme-suite.org/doc/centrimo.html).

### Analysis of publicly available data

Publicly available datasets (Moqtaderi et al. 2010; Oler and Cairns 2010; Sathira et al. 2010; Baillat et al. 2012; ENCODE 2012; Fong et al. 2017; Davis et al. 2018; Choquet et al. 2019) used in this study were listed in Table S3. Heatmap and average plot were generated using deepTools. For the analysis of pausing index, we defined proximal regions as −500 bp to +500 bp from the gene start site. Gene body regions were defined as +500 bp to the gene end, subtracted proximal regions of the internal gene start site of the isoforms.

### Data deposition

The data reported in this paper have been deposited in the Gene Expression Omnibus (GEO) database under accession number GSE133729.

## ACKNOWLEDGEMENTS

We thank A. Stutzman for help with the plasmid construction, and S. Pott, K. Ikegami, A. Ruthenburg, and members of Struhl laboratory for helpful comments and discussions. This work was supported by a fellowship from the University of Chicago Fellows Program to Y.H. and grants to K.S. from the National Institutes of Health (GM 30186 and CA 107486).

---

**Supplementary Fig. 1.**
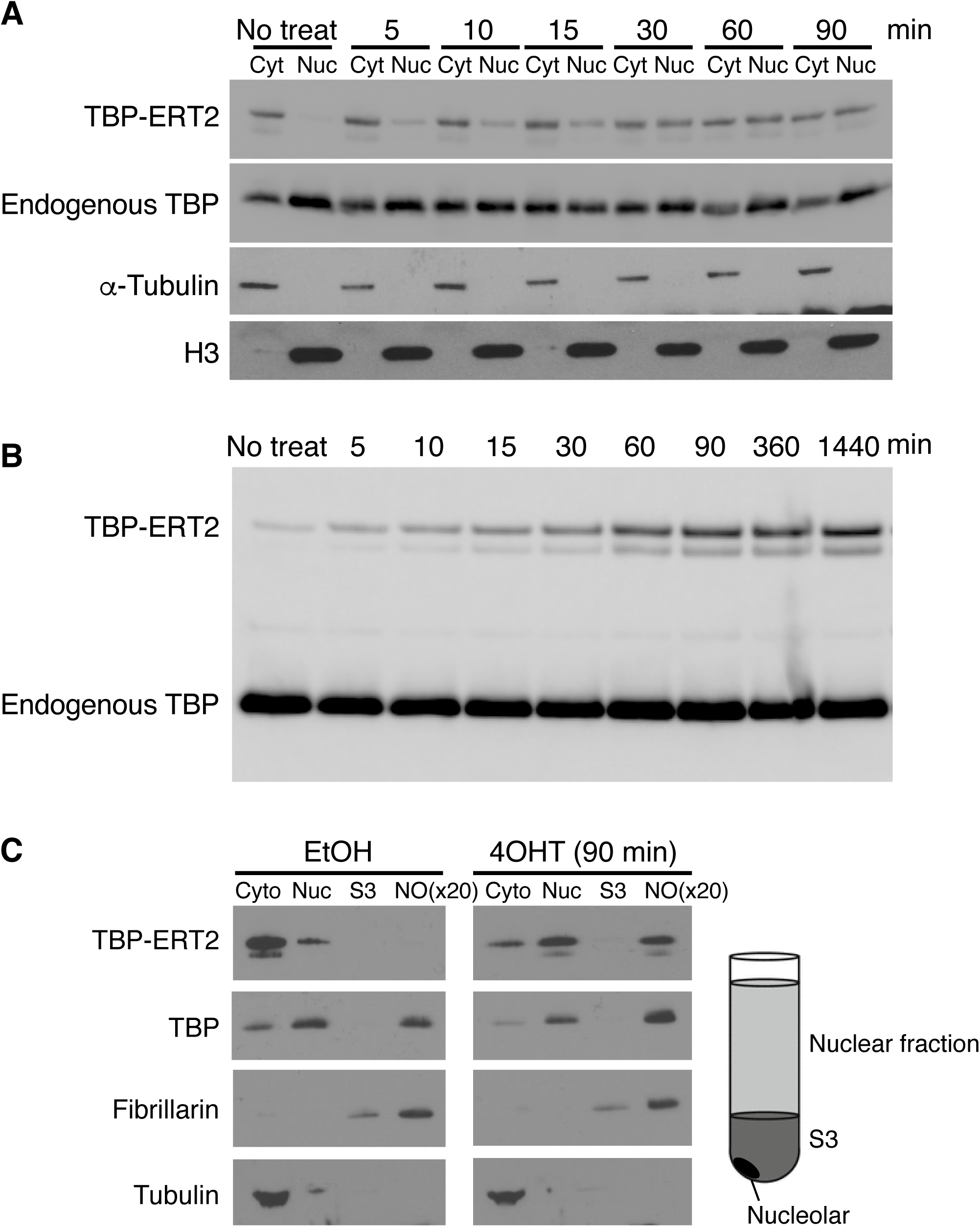
TBP-ERT2 localizes to the nucleus upon tamoxifen treatment. (*A*) Western blotting analysis of TBP-ERT2, endogenous TBP, *α*-Tubulin (cytoplasmic marker), and Histone H3 (nuclear marker) in the cytoplasmic (Cyt) and the nuclear (Nuc) fraction following the given time after tamoxifen treatment. (*B*) Western blotting analysis of TBP-ERT2 and endogenous TBP in nuclear fraction. (*C*) Western blotting analysis of TBP-ERT2, endogenous TBP, Fibrillarin (nucleolar marker), and *α*-Tubulin (cytoplasmic marker) in the cytoplasmic (Cyt), nuclear (Nuc), S3, and nucleolar (NO) fraction before (EtOH) and 90 min after (4OHT) tamoxifen treatment. Nucleolar fraction was concentrated to 20-fold.

**Supplementary Fig. 2.**
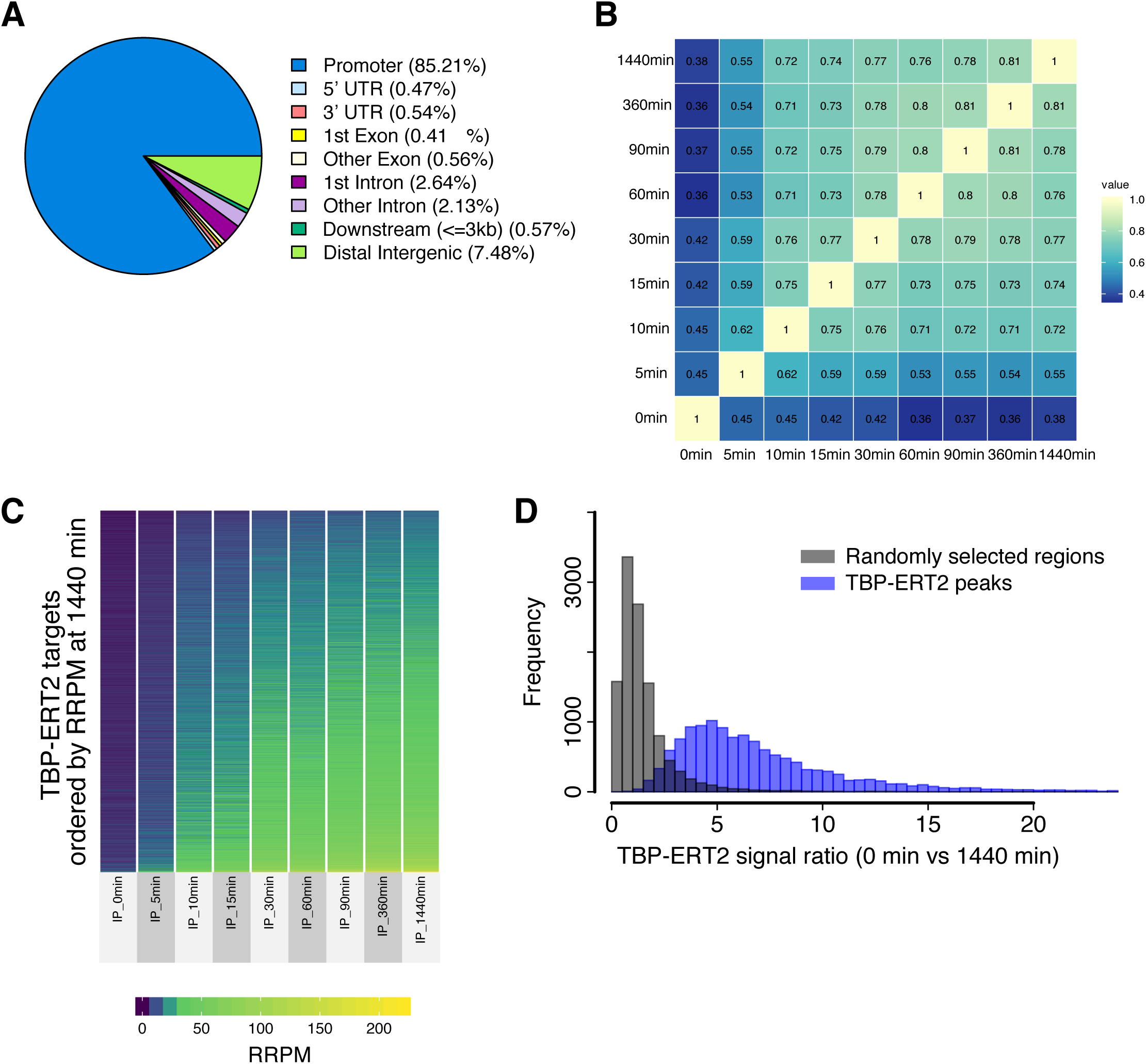
TBP-ERT2 ChIP signal gradually increases after tamoxifen treatment. (*A*) Location of TBP-ERT2 peaks. (*B*) Spearman correlation of ChIP-seq signal at peaks between samples at a given time point after tamoxifen induction. (*C*) ChIP-seq signal (RRPM: Reference-adjusted RPM) at each target site through time-course. (*D*) Histogram of the frequency (number) of regions or TBP-ERT2 peaks having indicated ChIP-seq signal ratio between 0 min and 1440 min samples.

**Supplementary Fig. 3.**
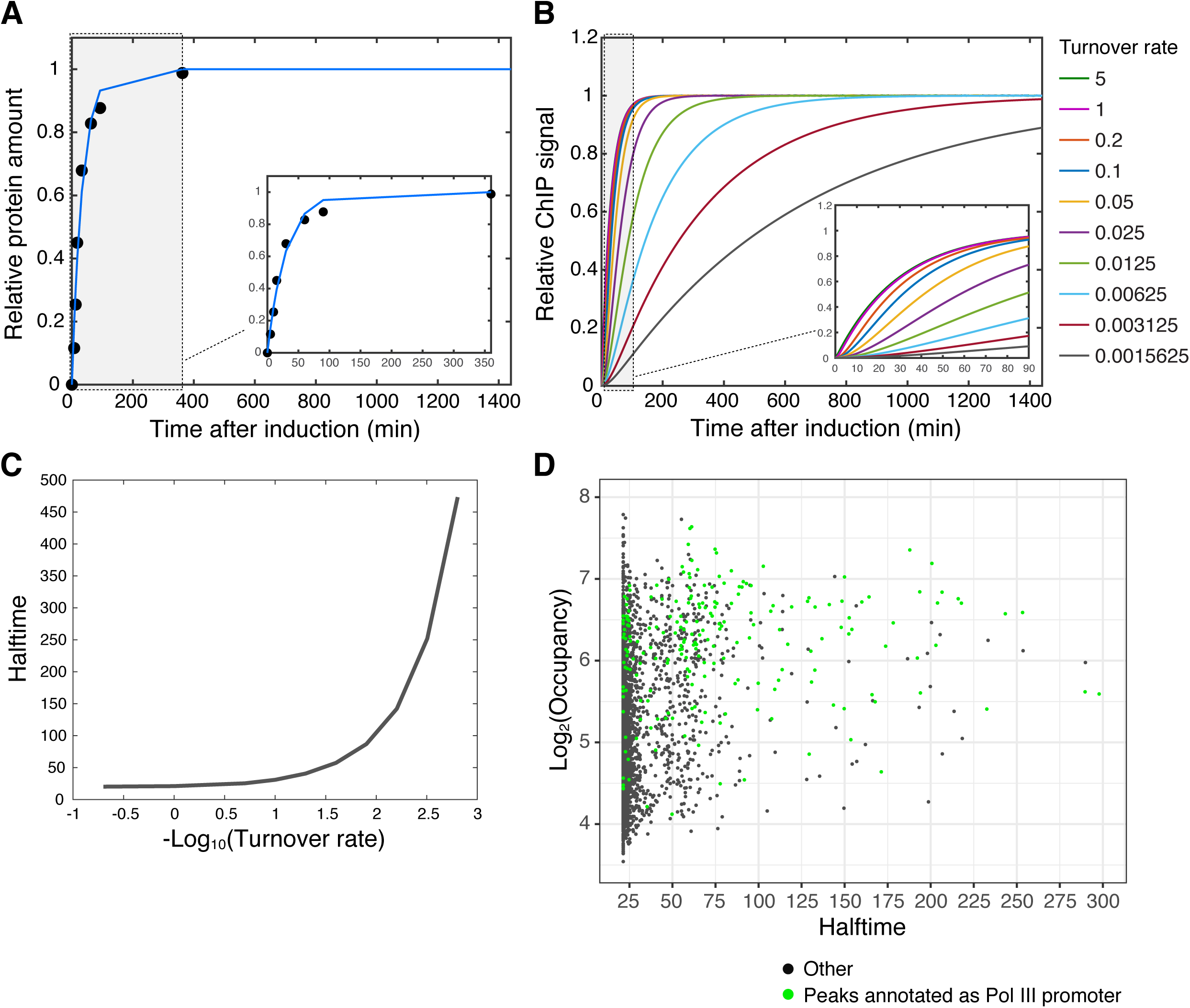
ChIP signal increasing is reflected in the binding turnover rate. (*A*) Change of the nuclear TBP-ERT2 amount over the time-course. Markers are real data measured by western blotting (means of the three independent analyses) and the line is simulated change of the protein amount based on the equation describing nuclear translocation of the protein (Methods). Small window indicates the enlargement of the data at the early time points. (*B*) Simulated increase of the relative ChIP signal with various Lambda (binding turnover rate) values. Small window indicates the enlargement of the data at the early time points. (*C*) Relation between Lambda (binding turnover rate) and halftime (time point where the relative ChIP signal reaches to 0.5). (*D*) Scatter plot of TBP-ERT2 peaks in log2 occupancy vs halftime. Blue color indicate peaks annotated as pol III genes. Black are the others.

**Supplementary Fig. 4.**
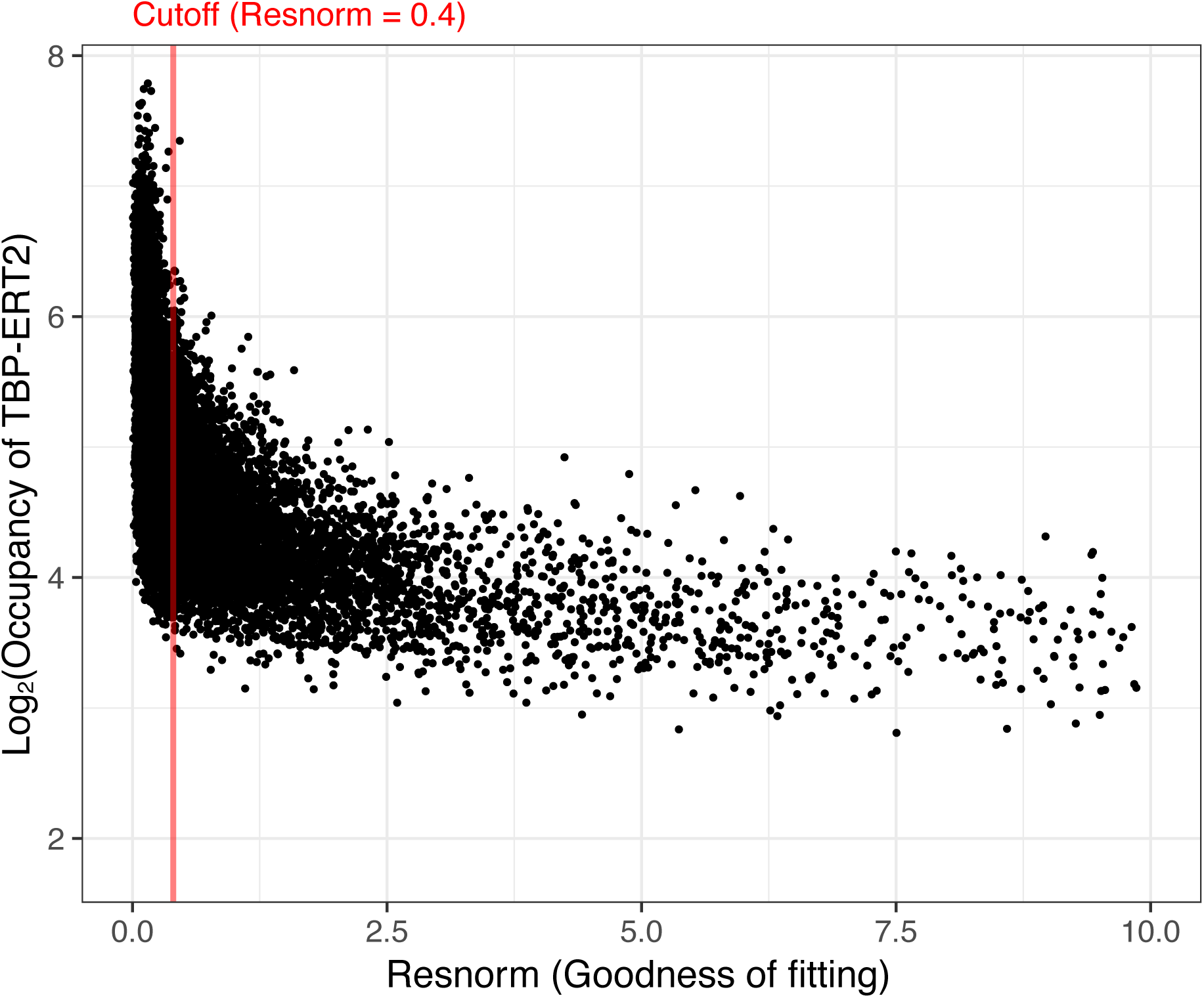
Cutoff for peaks that we set based on Goodness-of-fitting.

**Supplementary Fig. 5.**
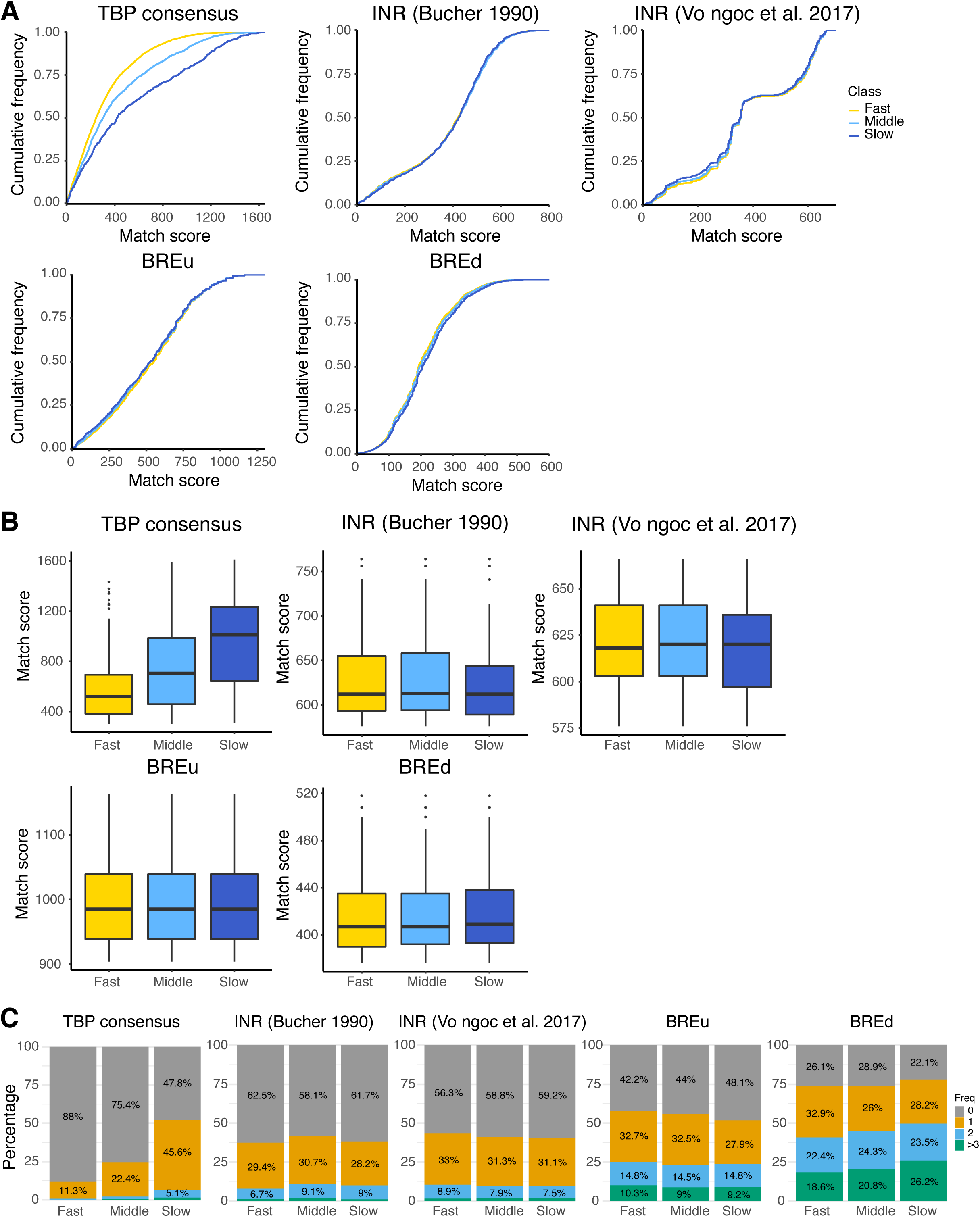
Analysis of motifs in Pol II core promoter. (*A*) Cumulative frequencies of match score of hit sequences to each motif in +/- 100 bp region from gene start. (*B*) Boxplots of the highest match score of hit sequences (forward direction) in each of the +/- 100 bp region from gene start. (*C*) Percentages of the +/- 100 bp region (Fast n=4,096, Middle n=1,115, Slow=412) that have hit sequences over the match score thresholds (corresponding to cut-off extracting approximately upper 25% of hit sequences: TBP consensus > 400, INR (Bucher 1990) > 575, INR (Vo ngoc, et al. 2017) > 625, BREu > 750, BREd > 300).

**Supplementary Fig. 6.**
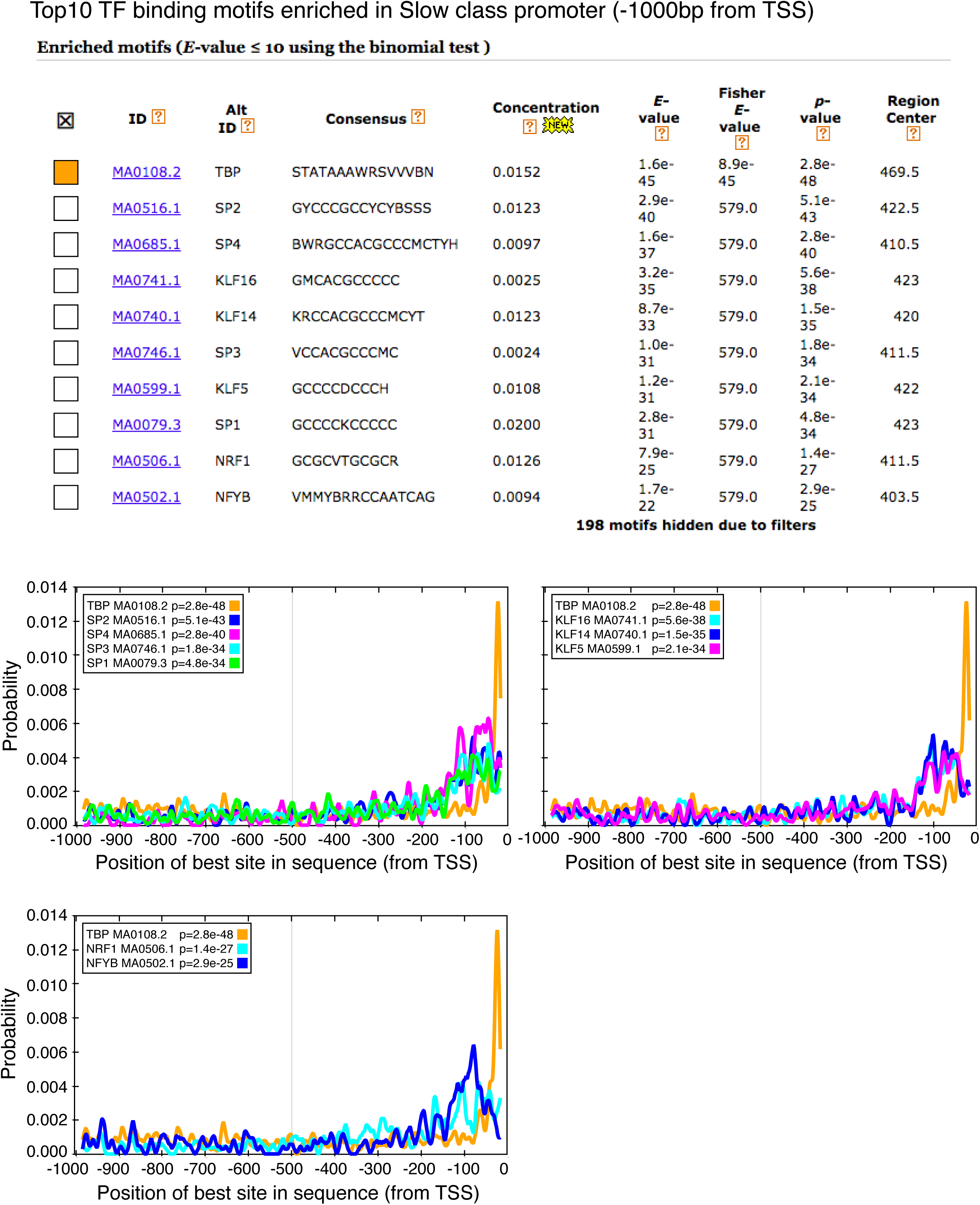
Known TF binding motifs enriched in slow class. Known TF binding motifs enriched in slow class searched by CentriMo (http://meme-suite.org/doc/centrimo.html).

**Supplementary Fig. 7.**
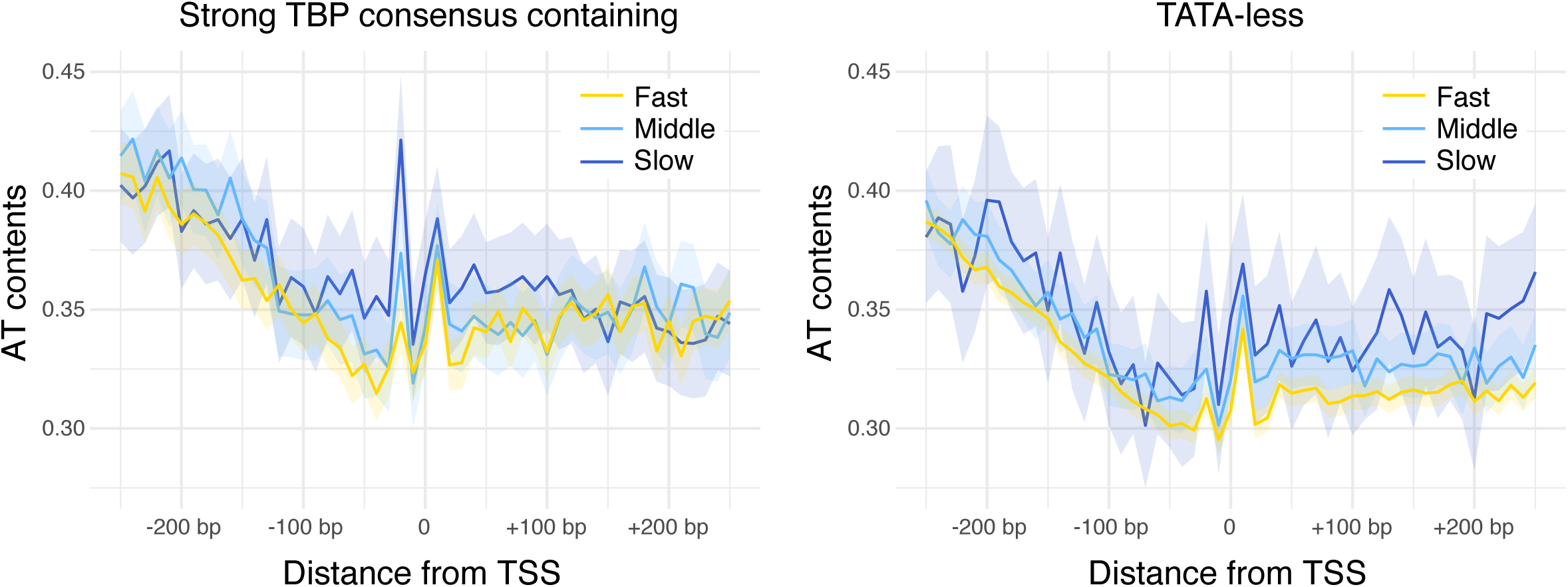
TATA-less pol II promoters in Slow class tend to have higher AT content. Average AT-contents in the promoters containing strong TBP consensus (left) or TATA-less promoters (right). Promoters that have strong TBP consensus (pval < 0.0001, PWMtools) or TATAWAW (RSAT, pattern matching) are categorized as Strong TBP consensus containing (n=1459, Fast n=895, Middle n=415, Slow n=149). Rest were categorized in TATA-less (n=4164, Fast n=3201, Middle n=700, Slow n=263). Shadow areas represent 95% confidence intervals for the means.

**Supplementary Fig. 8.**
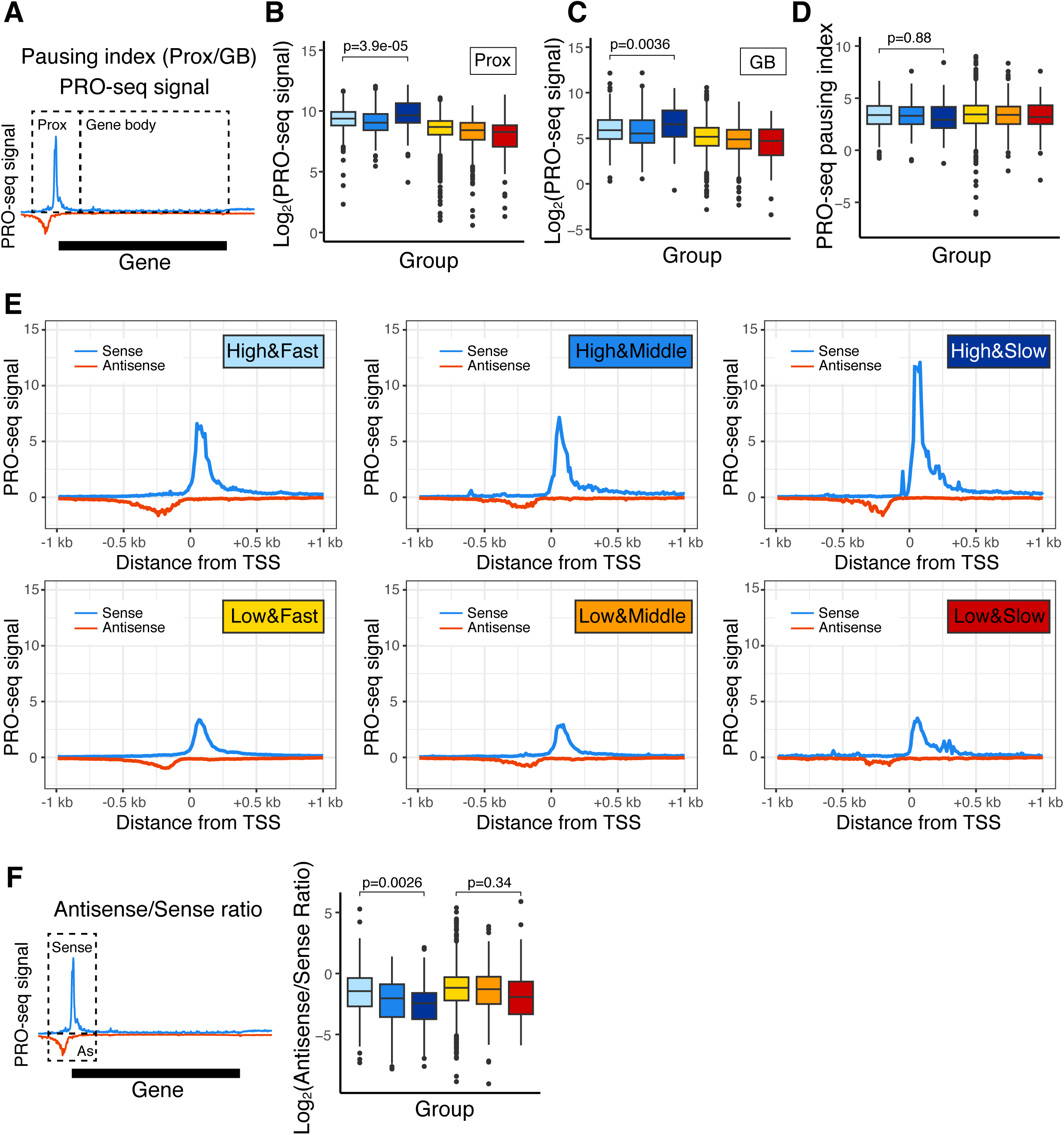
Polymerase pausing does not associate with the binding turnover rate of TBP, and high and slow class Pol II promoters show strong transcription in sense direction. (*A*) Schematic illustration of the proximal region (Prox: from −500-bp to +500-bp of TSS) and the gene body region (GB: from +500-bp of TSS to TES). (*B*) PRO-seq signal in the proximal region. (*C*) PRO-seq signal in the gene body (GB) region. (*D*) Pausing index. P-values are t-test results. (*E*) PRO-seq signal in sense (blue) and anti-sense (red) direction. (*F*) Schematic illustration of the region (from −500-bp to +500-bp of TSS) for counting PRO-seq signal (left), and log2 ratio between anti-sense and sense PRO-seq signal (right). P-values are t-test results.

**Supplementary Fig. 9.**
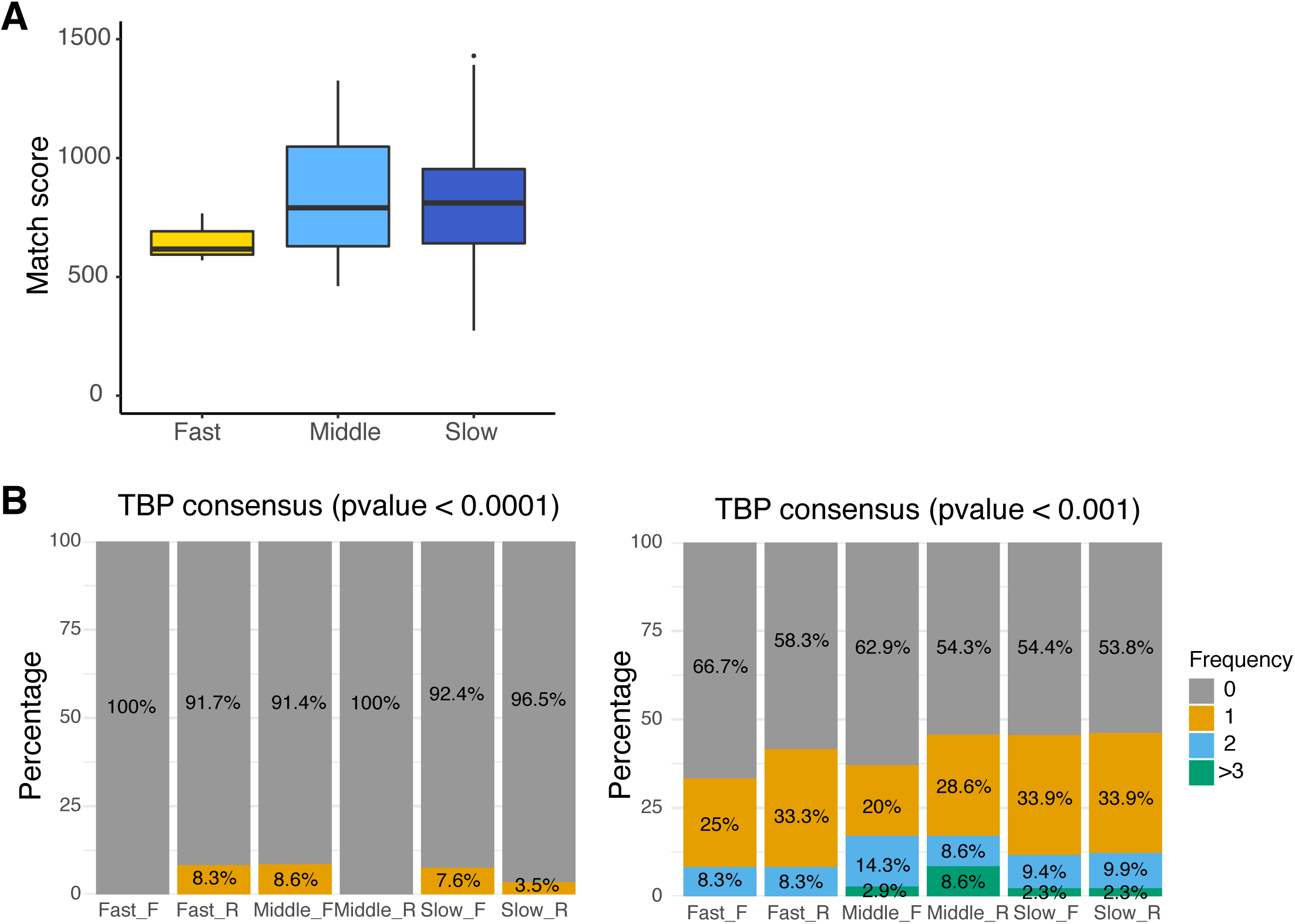
TBP consensus motif in pol III genes. (*A*) Boxplots of the highest match score of hit sequences in forward direction. (*B*) Percentages of the pol III promoters (pseudogenes were removed, n=217, Fast n=11, Middle n=35, Slow=171) that have hit sequences to TBP consensus motif (left: p-value < 0.0001, right: p-value < 0.001, PWMtools).

**Supplementary Table 1. Numbers of reads mapped and passed the quality filter**

**Supplementary Table 2. Primer list**

**Supplementary Table 3. GEO accession numbers of publicly available datasets used in this study**

